# Genetic Diversity of *Zymoseptoria tritici* Populations in Central and South-eastern Ethiopia

**DOI:** 10.1101/2024.10.02.616306

**Authors:** Ayantu Tucho, Tilahun Mekonnen, Farideh Ghadamgahi, Samrat Gosh, Diriba Muleta, Kassahun Tesfaye, Eu Shang Wang, Tesfaye Alemu, Ramesh Raju Vetukuri

**Affiliations:** Institute of Biotechnology, Addis Ababa University, College of Natural and Computational Sciences, P. O. Box 1176, Addis Ababa, Ethiopia, http://www.aau.edu.et/; Department of Plant Science, Salale University, P. O. Box 245, Fitche, Ethiopia; Department of Plant Breeding, Swedish University of Agricultural Sciences, SE-234 22 Lomma, Sweden; Department of Microbial, Cellular and Molecular Biology, P. O. Box 1176, Addis Ababa University, Ethiopia University; Bio and Emerging Technology Institute (BETin), P.O.Box:- 5954 Addis Ababa, Ethiopia

**Keywords:** AMOVA, Gene flow, Genetic analysis, ITS rDNA, SSR, Structure, *Z. tritici*

## Abstract

Septoria tritici blotch (STB), caused by the hemibiotrophic fungus *Zymoseptoria tritici*, is a serious threat to global wheat production, and a major bottleneck to wheat production in Ethiopia. Accurate identification and analysis of the pathogen’s genetic structure helps inform robust STB management. This study analyzed the molecular identity and genetic structure of 200 *Z. tritici* isolates retrieved from diseased crops in central and south-eastern regions of Ethiopia. Allelic diversity at 12 simple sequence repeat (SSR) loci was examined for all 200 isolates, and 165 isolates were identified by Sanger sequencing of the internal transcribed spacer (ITS) region of nuclear DNA (rDNA) region. The microsatellites were highly polymorphic, with mean number of alleles (Na) = 6.23, effective alleles (Ne) =2.90, Nei’s gene diversity (H) =0.57, and polymorphic information content (PIC) =0.59, and an analysis of molecular variance confirmed the presence of low population differentiation (Fixation Index (F_ST_) = 0.02, Gene Flow (Nm) = 14.7), with 95% of the total genetic variation residing within populations, and only 5% residing between populations. The highest genetic diversity (Number of allele (Na) = 9.33, Effective number of allele (Ne) = 3.41 and Nei’s gene diversity (H) = 0.68) was observed in the Oromia special zone surrounding Finfinnee (OSZ) *Z. tritici* populations, followed by Arsi and North Shewa populations, indicating that these areas are ideal for multi-location germplasm resistance screening, and also pathogen genetic and genomic analyses. Cluster analyses did not clearly divide the populations into genetically separate clusters according to their geographic sampling areas, probably because of high gene flow. All individual samples showed genetic admixture, and shared genomic backgrounds from two subgroups (K=2). Overall, the SSR markers proved to be highly informative and effective genetic tools for unlocking the pathogen’s genetic structure. The *Z. tritici* populations of central and southeast Ethiopia exhibit high genetic diversity, indicating the need to deploy durable and diverse disease management strategies. North Shewa, OSZ, Arsi and West Arsi administrative zones represent hotspots for genetic and genomic analyses of *Z. tritici*. Moreover, these areas would be excellent locations for host–pathogen interaction studies, and wheat germplasm screening for STB resistance.

## 1 Introduction

*Zymoseptoria tritici* (*Mycosphaerella graminicola*), the causal agent of septoria tritici blotch (STB), which affects bread wheat (*Triticum aestivum* L.) (McDonald *et al*., 2015), is a hemibiotrophic apoplastic fungal species (Ma *et al*., 2018). It belongs to the class Ascomycete (Testa *et al*., 2015), and has a haploid genome of 21 chromosomes: 13 core chromosomes and 8 accessory chromosomes (Goodwin *et al*., 2011). *Z. tritici* possesses a heterothallic mating system, which requires two compatible partners of opposing mating types (mat1-1 and mat1-2) to come together to create sexual spores (Waalwijk *et al*., 2002). STB is the most destructive disease of wheat across the world, causing yield losses of up to 50% in untreated fields planted with susceptible cultivars (Eyal and Levy, 1987, McDonald *et al*., 2015). Primary infections of STB can occur from airborne or rain-splashed asexual pycnidospores and sexual ascospores from infected crop debris (Hunter *et al*., 1999).

Bread wheat is one of the most important global cereals, supporting nearly 35% of the world’s population as a staple food (statista, 2023). South Africa and Ethiopia are the largest wheat producers in Sub-Saharan Africa (Dibaba, 2019). In Ethiopia, wheat ranks fourth after teff (*Eragrostis tef*), maize (*Zea mays*) and sorghum (*Sorghum bicolor*) in area coverage, and third after maize and teff in total production (Bezabih *et al*., 2023). It is cultivated by nearly 5 million householders, on about 1.7 million ha, for various purposes, including as food, animal feed and income generation (Latta *et al*., 2013). Bread wheat accounts for nearly 80% of the total wheat production of the country (Latta *et al*., 2013, Bezabeh *et al*., 2014). In 2022, 2.3 million ha of Ethiopia was covered with wheat, with a total production and productivity of 7 million metric t and 3.04 t/ha, respectively (FAOSTAT, 2024). This national average productivity (3.04 t/ha) is far lower than the global average of 3.69 t/ha (FAOSTAT, 2024) and the productivity level in USA (3.5 t/ha), resulting in a production limit to meeting the growing demand for food by the ever-increasing population of the country (Gemechu and Tadese, 2018). Limited access to advanced production technologies, low agricultural inputs (fungicides, improved varieties and fertilizer), and biotic and abiotic stresses are some of the major wheat production constraints in Ethiopia.

In Ethiopia, wheat cultivation is persistently affected by over 30 fungal diseases (Bekele, 1985), of which stripe (yellow) rust caused by *Puccinia striiformis* f. sp. *tritici* (Pst), stem rust caused by *P. graminis* f. sp. *tritici*), leaf rust caused by *P. triticina*, and STB caused by *Z. tritici* are the most destructive (Huluka, 2002, Hailu *et al*., 2015, Mann and Warner, 2017). Rusts can result in grain yield losses of 60–100% (Olivera *et al*., 2015). About 25–82% of wheat production loss has been documented as caused by STB, and the disease incidence and severity have increased recently in Ethiopia’s major wheat belts (Mekonnen et al., 2020). STB infestation results in significant grain yield loss by reducing tillering, poor seed set, poor grain fill or shriveled kernels, and death of leaves, spikes or the entire plant (Mekonnen *et al*., 2020, Goodwin *et al*., 2011).

The narrow genetic basis of modern wheat cultivars and rapidly changing fungal genomes may have resulted in the frequent breakdown of host resistance (Ye *et al*., 2019). The sexual cycle of *Z. tritici*, synonyms *M. graminicola* plays a crucial role in its epidemiology and genetic diversity among field isolates. Resistance breeding provides the best eco-friendly approach to managing STB. However, while resistance breeding contributes to the development of cultivars that are durable and have broad-spectrum resistance to diseases, pests and weeds, and enables screening for locally adapted varieties, understanding the pathogen’s genetic diversity and population structure is crucial for implementing proper control measures and developing novel management strategies (Siah *et al*., 2018). Knowledge of the pathogen’s population structure can enhance predictions of the effectiveness and durability of host resistance. Plant pathogen populations with high genetic variability are more likely to adapt to resistant cultivars than populations with low genetic variability. Understanding the evolutionary forces controlling pathogen populations therefore holds enormous potential for the development and implementation of robust disease management measures (McDonald *et al*., 1995, McDonald *et al*., 2015).

The advent of molecular marker technology has promising applications in plant pathology, in assessing genetic variation among pathogens (Ramesh *et al*., 2020) and fingerprinting pathogen ecology and epidemiology. Internal transcribed spacer (ITS) sequences within ribosomal RNA genes (rDNA) are widely used as molecular probes to identify fungal pathogens (Yang *et al*., 2018). It is the prime fungal default region for species identification, and it can be retrieved using diagnostic primers that target the sequence (Das and Deb., 2015). However, research efforts to identify *Z. tritici* populations in Ethiopia based on ITS region sequencing are greatly lacking. Moreover, information on the genetic structure of the pathogen populations in the central and south-eastern parts of the country is limited. DNA markers such as simple sequence repeats (SSR) have been widely used to disclose the genetic structure of *Z. tritici* populations in various countries, such as Tunisia (Boukef *et al*., 2013), England (Owen *et al*., 1998) and France (El Chartouni *et al*., 2011a);(Siah. *et al*., 2018). Therefore, the aim of this study was the molecular identification and genetic analysis of *Z. tritici* populations recovered from central and south-eastern regions of Ethiopia.

## 2 Materials and Methods

### 2.1 Study areas and sampling procedure

*Z. tritici* isolates were recovered from STB-infected wheat leaf samples collected from six major wheat-growing zones in Ethiopia: West Shewa (WSH), Southwest Shewa (SWSH), North Shewa (NSH), Oromia special zone surrounding Finfinnee (OSZ), West Arsi (WA) and Arsi zone (AZ). These regions together represented 75% of the country’s total wheat production area. Samples were collected during the main cropping seasons (summer) of 2021, by following the main roads and accessible routes in a randomly selected major wheat-growing district. Stops were made at 5 – 10 km intervals where wheat fields were accessible, based on vehicle odometers. Figure 1 and Table 1 provide a map of the sample collection sites and summary information regarding the six *Z. tritici* populations considered. During collection, green leaves with symptoms of STB were collected in paper envelopes labelled with the sample code. The collected samples were then air-dried at room temperature and stored at 5°C until isolation.

**Figure 1.**
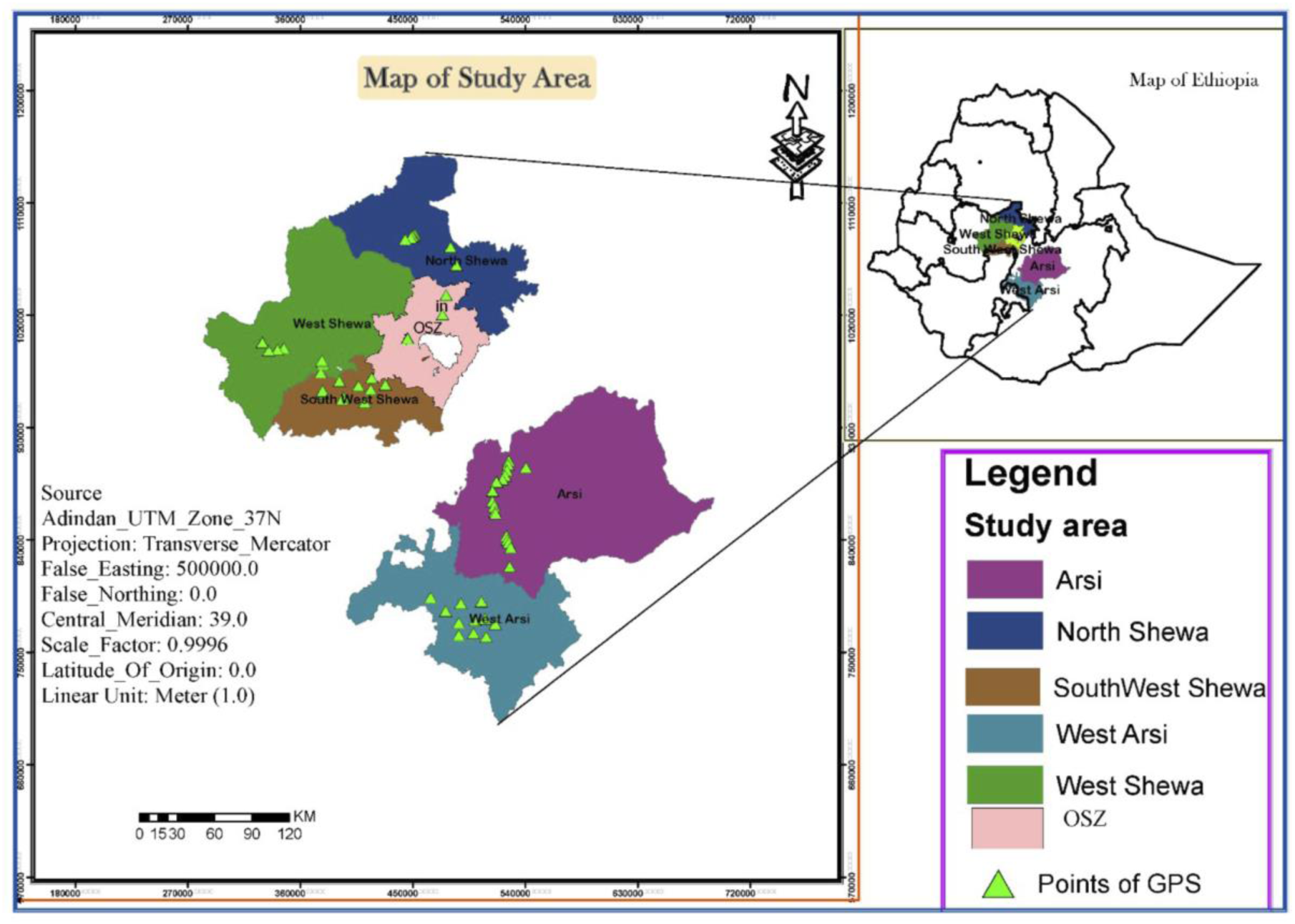
A map of Ethiopia showing the six administrative zones used for the collection of *Zymoseptoria tritici* isolates. GPS = global positioning system, OSZ = Oromia special Zone Surrounding Finfinne.

**Table 1.** Summary information for the six populations of *Zymoseptoria tritici* sampled in Ethiopia.

| Population | Isolate code | Geographical position (UTM) |  | Elevation (m) |
| --- | --- | --- | --- | --- |
|  |  | Latitude range | Longitude range |  |
| Arsi | ZTET043 – ZTET119 | 07° 47' 775–08°17' 474 | 039°12' 465–039°26' 206 | 1672–2856 |
| North Shewa | ZTET179 – ZTET200 | 09°58' 761–09°81' 214 | 038°48' 647–038°86' 122 | 2550–3101 |
| SW Shewa | ZTET145 – ZTET160 | 08°617'289–08°74'708 | 037°87'918–038°04'177 | 2259–2853 |
| West Arsi | ZTET120 – ZTET144 | 07°01'438–08°12'959 | 038°78'611–039°36'766 | 1994–2713 |
| West Shewa | ZTET161 – ZTET178 | 08°81'051–09°02'660 | 037°44'785–037°89'241 | 2230–3366 |
| OSZ | ZTET001 – ZTET042 | 08°88'092–09°37'329 | 038°50'680–038°81'904 | 2158–2604 |

UTM = Universal Transverse Mercator coordinate system. SW Shewa = Southwest Shewa; OSZ =
Oromia special zone surrounding Finfinnee.

### 2.2 Fungal isolation procedures

Spore isolation and subsequent laboratory activities were carried out at the National Agricultural Biotechnology Research Centre (NABRC), Holeta, located 29 km west of Addis Abeba, Ethiopia. Fungal isolation was conducted following the procedure of (Eyal and Levy, 1987), with some modifications. Symptomatic leaf samples were cut into about 10-cm lengths and placed on sterile filter paper in Petri dishes wetted with sterile distilled water. The Petri dishes were then placed in polyethylene plastic bags and incubated for 3–4 h at 24°C. The samples were periodically checked under a stereoscopic dissecting microscope for the formation of a cloudy ooze on the top of the pycnidia. Using a flame-sterilized fine needle, the mono-pycnidia oozing drops were transferred onto potato dextrose agar (PDA; potato 200 g/l, dextrose 20 g/l, agar 15 g/l). Colony morphology was confirmed under a microscope (40× objective, using a Konsortiet Laborlux 11 microscope; Leitz Wetzlar, Vienna, Germany), and creamy pink-colored colonies were streaked onto new PDA plates supplemented with 250 mg chloramphenicol. Inoculated Petri dishes were kept at 24°C for 7–10 days until fungal growth was observed. Single spore-derived colonies were transferred into a liquid medium composed of 1% (w/v) yeast extract powder + 1% (w/v) sucrose, and the cultures were maintained on an orbital shaker at 130 rpm for 3 weeks, for spore multiplication. The fungal spores were then collected in Eppendorf tubes for molecular diversity analysis. In total, 200 single-spore-derived isolates were successfully retrieved and preserved in 25% glycerol at –80°C for further study.

### 2.3 Molecular identification of *Z. tritici* isolates

DNA extraction was carried out at the Swedish University of Agricultural Science (SLU), Department of Plant Breeding, at Alnarp, Sweden. The glycerol-preserved cultures were shipped to SLU and transferred on arrival to PDA (Sigma-Aldrich, Hamburg, Germany). Ten-day-old isolates grown on PDA were used for DNA isolation. DNA extraction was done using a Quick-DNA^TM^ fungal/bacterial miniprep kit following the manufacturer’s protocol (Zymo Research, Irvine, CA, USA). The extracted DNA quality was checked by loading 5 µl DNA + 2 µl 6× loading dye with gel red on a 1% agarose gel and separated at 100 V for 40 min. The DNA concentration was checked using a Nano-drop spectrophotometer (DS-11 Series Spectrophotometer/Fluorometer; De Novix, Wilmington, DE, USA). The isolates were confirmed using a universal primer (ITS5 forward and ITS4 reverse) (White *et al*., 1990) (Table 2) that targeted ITS sequences of rDNA. A polymerase chain reaction (PCR) reaction was carried out in a 46-μl reaction volume composed of 22 μl phire^TM^ plant direct master mix, 1 μl each of the forward (ITS5) and reverse (ITS4) primers, 21 μl nuclease-free water and 1 μl genomic DNA, using a Bio-Rad Thermal cycler PCR (Bio-Rad, Hercules, CA, USA). The PCR program followed an initial denaturation at 98°C for 5 min followed by 40 cycles of 5 s denaturation at 98°C, 1 min annealing at 55°C, primer extension at 72°C for 1 min, and a final extension step of 10 min at 72°C. The PCR product was run on 2% agarose gel electrophoresis. A 1 kb marker was used to estimate the product size. The PCR products were then purified using a Quick-DNA™ fungal/bacterial miniprep purification kit following the manufacturer’s protocol. The purified product quality was checked using Nano-drop spectrophotometers (DS-11 Series Spectrophotometer/Fluorometer; De Novix, Wilmington, DE, USA) and sent for Sanger sequencing (Eurofins Genomics, Ebensberg, Germany). The sequence results of 195 isolates were blasted [using the Basic Local Alignment Search Tool (BLAST), nih.gov] by using the website of National Center for Biotechnology Information (NCBI) and a phylogenetic tree was constructed by using Fast Tree (Price *et al*., 2009), then a Newick format tree annotated using ITOL v6 (Letunic and Bork, 2024).

**Table 2.**
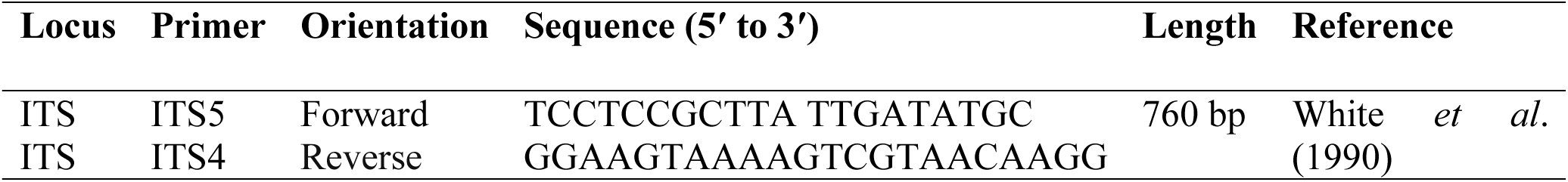
Primer sequences for amplification of the internal transcribed spacer (ITS) for the molecular identification of *Zymoseptoria tritici*.

### 2.4. Genetic structures using SSR markers

#### 2.4.1. Genotyping

The genetic structure of 200 isolates was explored using polymorphic published microsatellite markers. Twelve simple sequence repeat (SSR) markers (Table 3) were selected based on good amplification, polymorphism, specificity and suitability for multiplexing with DNA samples. The forward primers were 5′-labeled with four different fluorophore fluorescent dyes: FAM™, TAM™, ROX™ and HEX™ (Eurofins Genomics Ebensberg, Germany).

**Table 3.**
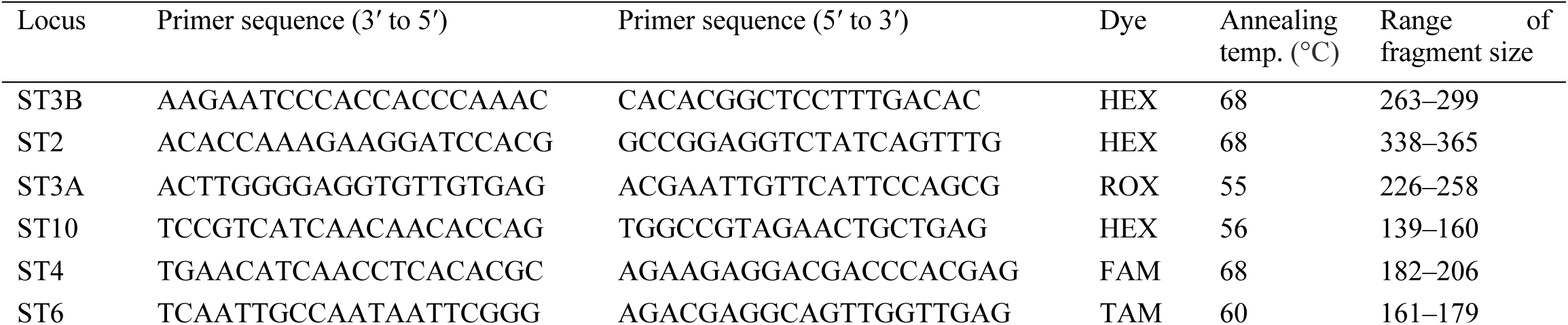
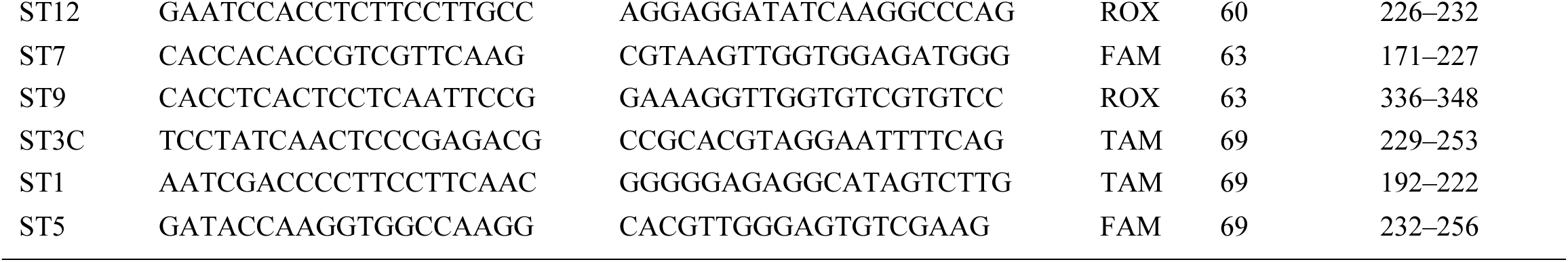
The 12 simple sequence repeat (SSR) markers used for genetic diversity analyses of *Zymoseptoria tritici*.

The PCR was performed in a total volume of 12.5 μl containing 6.75 μl phire™ plant direct master mix, 3.75 μl nuclease free water, 0.5 μl each of the forward and reverse primers, and 1 μl genomic DNA. A S1000™ Thermal Cycler (Bio-Rad) was used for amplification of the target loci. The primers’ annealing temperatures were gradient-optimized. The PCR program followed an initial denaturation at 95°C for 5 min followed by 35 cycles of denaturation at 95°C for 1 min, primer annealing at optimized temperatures of 55°C, 56°C, 60°C, 63°C and 69°C for 1 min (Table 3), and primer extension at 72°C for 2 min, followed by a final extension at 72°C for 10 min and holding at 4°C. PCR amplification was verified before capillary electrophoresis by running 5 µl of the products on a 1.5% agarose gel with Gel Red and visualizing the gel using a BioDoc-It™ Imaging System (Upland, CA, USA). Twelve primer pairs of PCR products were multiplexed into three panels, each of which held four PCR products. To prevent overlapping, there had to be at least an 80 bp size difference between the PCR products in each panel that was labelled with the same fluorescent dye. The capillary electrophoresis of the PCR products was performed denatured in form amide at 99°C for 3 min, and analyzed using the SeqStudio™ 8 Flex Genetic Analyzer (Thermo Fisher Scientific, Waltham, MA, USA) at the Department of Plant Breeding, Swedish University of Agricultural Sciences (Alnarp, Sweden).

#### 2.4.2. Data analysis

Following capillary electrophoresis, peak identification at the suggested threshold intensity was conducted using the software Gene Marker version 3.0.1 (Soft Genetics, LLC, State College, Pennsylvania, USA) with default parameters. The GS600 size standard was used to determine the fragment size. Each peak was considered as an allele at a codominant locus, and each individual or pool genotype at each locus was noted. Power marker version 3.25 (Liu and Muse, 2005) was used to calculate the locus diversity parameters, including polymorphic information content (PIC), major allele frequency (MAF), observed heterozygosity (Ho) and gene diversity [expected heterozygosity (He)] throughout the entire population. The software GenAlEx version 6.501 (Peakall and Smouse, 2012) was used to compute Nei’s gene diversity (H), the effective number of alleles (Ne), Shannon’s information index (I), allelic frequency and other population diversity indices, including the number of alleles (Na), the number of private alleles (NPA), and the percentage of polymorphic loci (PPL) across all loci for each population. A genetic differentiation test (PhiPT, with p-values across 999 bootstrap replications) and pairwise population genetic distances and gene flow were analyzed using the same software. The mafft aligner was used for multiple sequence alignment (Yamada *et al*., 2016), and Fast Tree was employed to construct the tree. The constructed Newick format tree was annotated using ITOL version 6 (Letunic and Bork, 2024). R package gg fortify was used for the principal component analysis (PCA). Arlequin version 3.5.2.2 was used to compute the analysis of molecular variance (AMOVA) and estimate the variance components (Excoffier and Lischer, 2010). R scripts within the gg plot R statistical package were used to generate a heatmap of pairwise Fixation Index (F_ST_) using the resulting sequences. The equation Nm (Haploid) = [(1/PhiPT) – 1]/2 was used to calculate the gene flow (Nm) across populations, where PhiPT stands for the variance between populations/total genetic variants. A genetic dissimilarity matrix was created (Nei, 1972) based on Neighbour Joining (NJ) and Nei’s standard genetic distance (DST, corrected), as well as the continuous Euclidian dissimilarity index.

The software STRUCTURE version 2.3.4 was used with a Bayesian model-based clustering technique to analyze the population structure and admixture trends (Pritchard *et al*., 2000). Data was gathered over 250,000 Markov Chain Monte Carlo (MCMC) replications for Population Cluster (K) = 1–10 using 20 iterations for each K, with a burn-in period of 100,000 utilized in each run to estimate the true number of population clusters (K). The web-based STRUCTURE HARVESTER version 0.6.92 was used to forecast the ideal K value using the simulation method developed by (Evanno *et al*., 2005). Using the Clumpak beta version, a bar plot for the ideal K was generated (Kopelman *et al*., 2015).

## 3 Results

### 3.1 Molecular identification of *Z. tritici*

Out of 200 *Z. tritici* isolates sent for Sanger sequencing, 167 of them maintained the quality control and BLAST-searched NCBI levels needed to confirm fungal identity. A phylogenetic tree was then generated (Figure 2), revealing high levels of genetic admixture. The isolates did not cluster according to the geographical areas of sampling.

**Figure 2.**
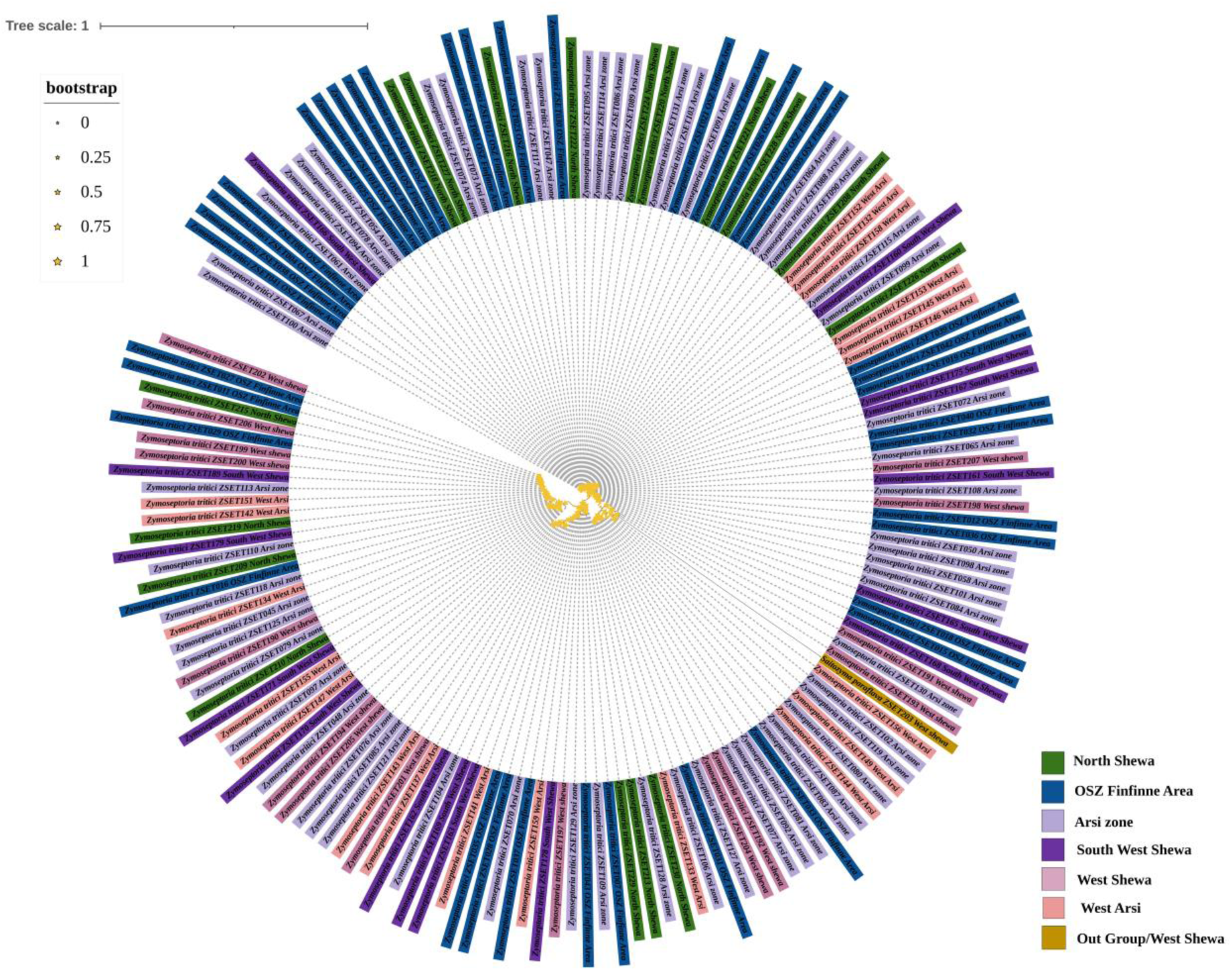
The phylogenetic tree generated for 167 *Zymoseptoria tritici* isolates collected from central and southeastern Ethiopia based on internal transcribed spacer (ITS) sequences within ribosomal RNA genes (rDNA). One isolate (*Saitozyma paraflava*) was Out of the Group which mean it is not *Zymoseptoria tritici* which is from west Shewa.

### 3.2 Genetic structure analyses

All 12 SSR loci were polymorphic (Figure 3 and Figure 4) and produced a total of 66 alleles with an average of 5.5 alleles per locus (Table 4). For each locus, the frequency of the most common allele was less than 0.95 or 0.99, confirming the high polymorphism of the markers. They were highly informative, with mean values for Na of 6.32 (range 3.83–10.83), Ne of 2.90 (range 1.90–4.89), I of 1.22 (range 0.78–1.8), H of 0.57 (range 0.44–0.73), genetic differentiation statistics by locus (Gst) of 0.01 (range 0.01–0.03) and Ho of 0.77 (range 0.64–0.91). The mean locus value for total expected heterozygosity (Ht) =0.81, Fixation index (F_ST_) =0.03 and Nei’s gene diversity (H) =0.62. (Table 4).

**Figure 3.**
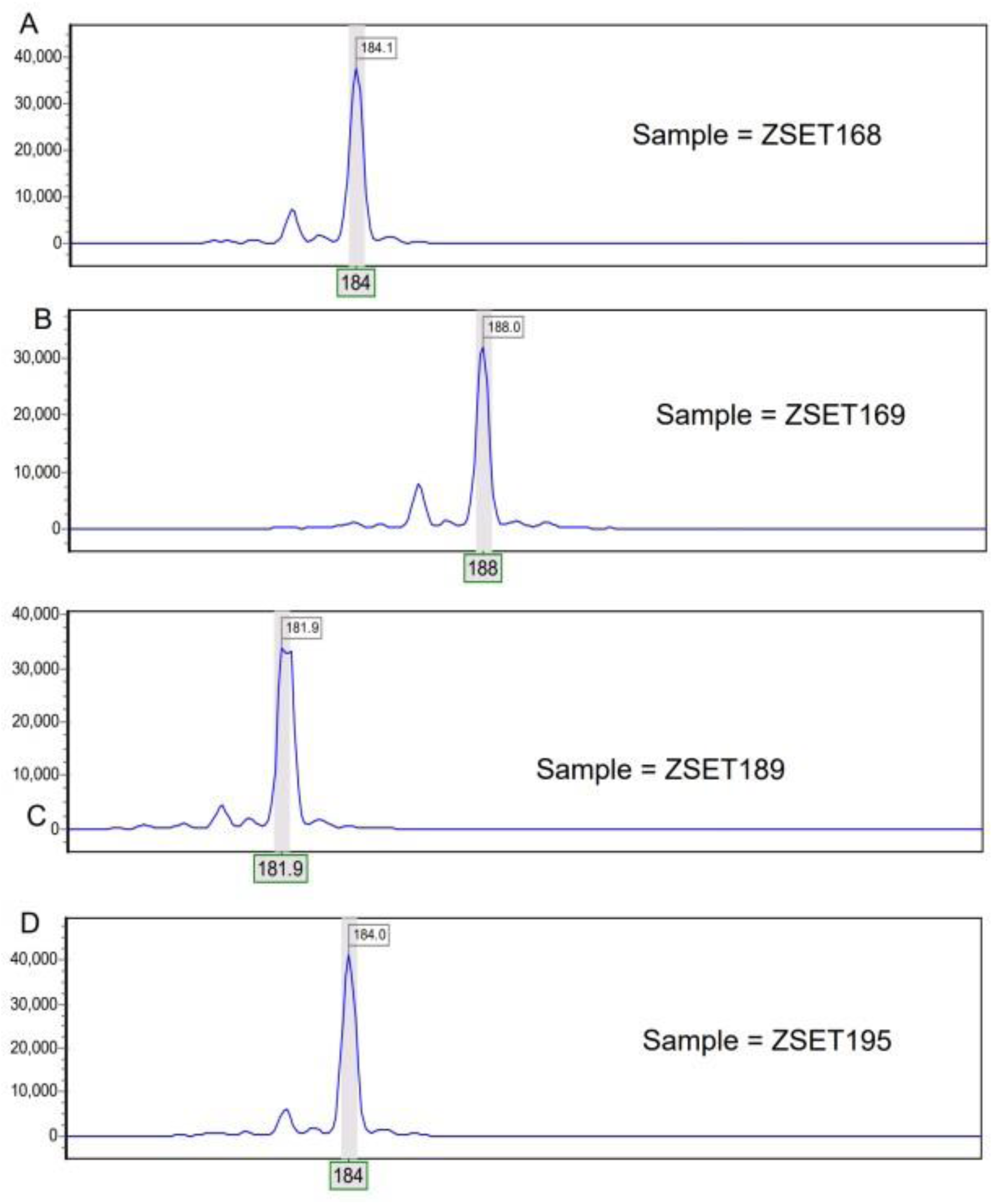
Capillary Electrophoresis result or Electrophoretograms for four *Zymoseptoria tritici* isolates at locus ST2 showing different sizes (bp) of amplification: **(A)** 184 bp in isolate ZSET168, **(B)** 188 bp in ZSET169, **(C)** 182 bp in ZSET189, and **(D)** 184 bp in ZSET168.

**Figure 4.**
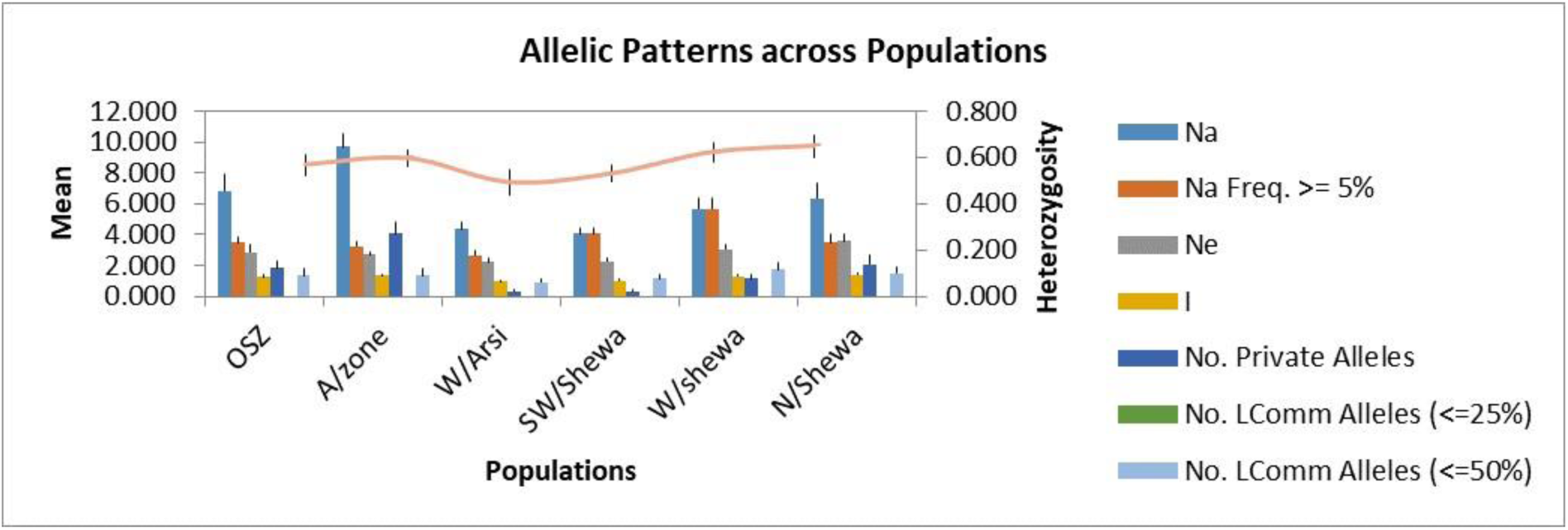
Allelic patterns across six populations of *Zymoseptoria tritici*. OSZ = Oromia special zone surrounding Finfinnee; A/Zone = Arsi; W/Arsi = West Arsi; SW/Shewa = Southwest Shewa; W/Shewa = West Shewa; N/Shewa = North Shewa. Na = numbers of different alleles; Na (Freq ≥5%) = numbers of different alleles with a frequency of ≥5%; Ne = numbers of effective alleles; I = Shannon’s Information Index; No. Private Alleles, numbers of alleles unique to a single population; No. LComm Alleles (≤25%) = numbers of locally common alleles (Freq. ≥ 5%) found in 25% or fewer populations and No. LComm Alleles (≤50%) = numbers of locally common alleles (Freq. ≥ 5%) found in 50% or fewer populations.

**Table 4.**
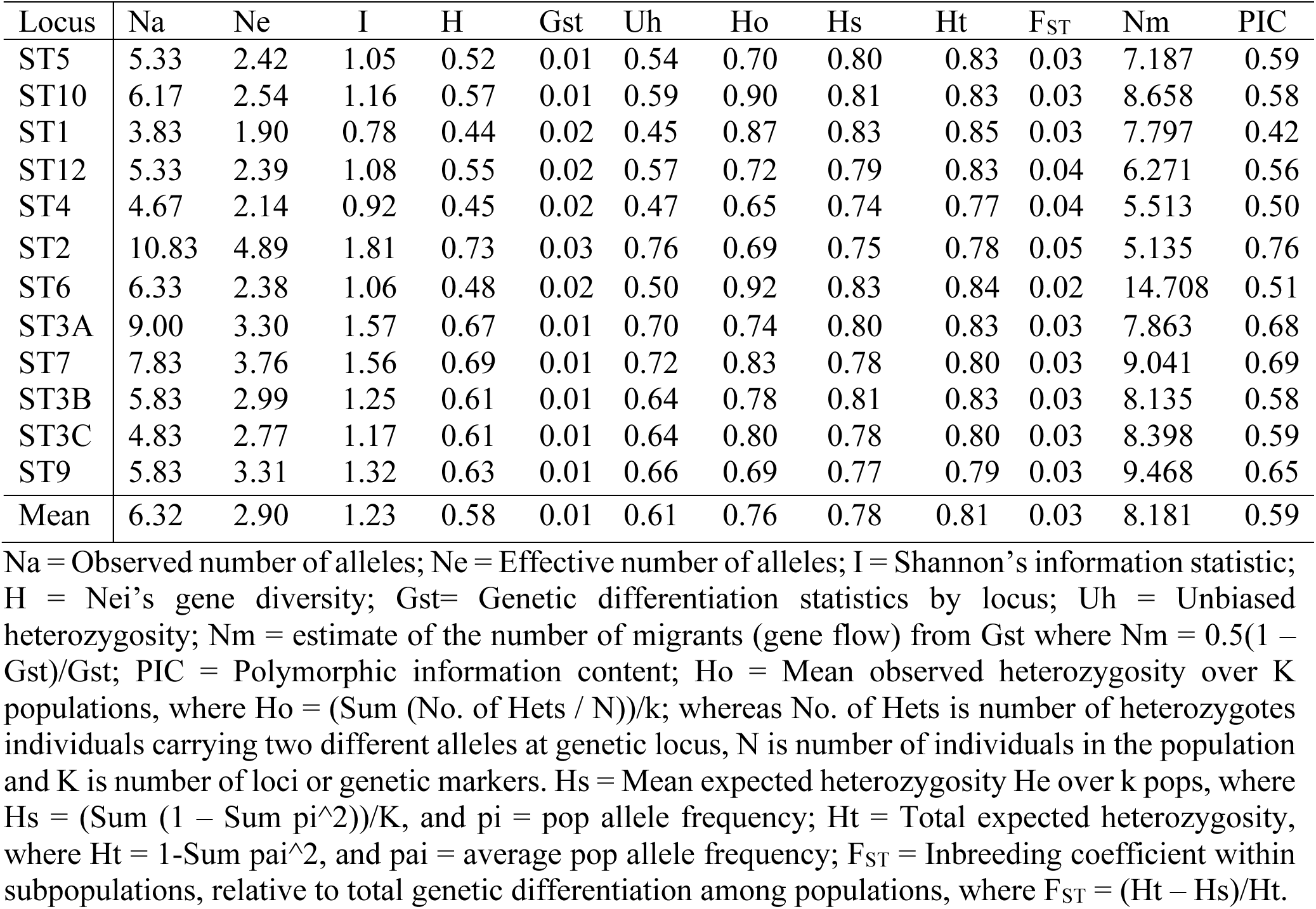
In formativeness and other genetic diversity summary statistics for 12 microsatellite loci across six populations of *Zymoseptoria tritici* in Ethiopia.

| Locus | Na | Ne | I | H | Gst | Uh | Ho | Hs | Ht | F<sub>ST</sub> | Nm | PIC |
| --- | --- | --- | --- | --- | --- | --- | --- | --- | --- | --- | --- | --- |
| ST5 | 5.33 | 2.42 | 1.05 | 0.52 | 0.01 | 0.54 | 0.70 | 0.80 | 0.83 | 0.03 | 7.187 | 0.59 |
| ST10 | 6.17 | 2.54 | 1.16 | 0.57 | 0.01 | 0.59 | 0.90 | 0.81 | 0.83 | 0.03 | 8.658 | 0.58 |
| ST1 | 3.83 | 1.90 | 0.78 | 0.44 | 0.02 | 0.45 | 0.87 | 0.83 | 0.85 | 0.03 | 7.797 | 0.42 |
| ST12 | 5.33 | 2.39 | 1.08 | 0.55 | 0.02 | 0.57 | 0.72 | 0.79 | 0.83 | 0.04 | 6.271 | 0.56 |
| ST4 | 4.67 | 2.14 | 0.92 | 0.45 | 0.02 | 0.47 | 0.65 | 0.74 | 0.77 | 0.04 | 5.513 | 0.50 |
| ST2 | 10.83 | 4.89 | 1.81 | 0.73 | 0.03 | 0.76 | 0.69 | 0.75 | 0.78 | 0.05 | 5.135 | 0.76 |
| ST6 | 6.33 | 2.38 | 1.06 | 0.48 | 0.02 | 0.50 | 0.92 | 0.83 | 0.84 | 0.02 | 14.708 | 0.51 |
| ST3A | 9.00 | 3.30 | 1.57 | 0.67 | 0.01 | 0.70 | 0.74 | 0.80 | 0.83 | 0.03 | 7.863 | 0.68 |
| ST7 | 7.83 | 3.76 | 1.56 | 0.69 | 0.01 | 0.72 | 0.83 | 0.78 | 0.80 | 0.03 | 9.041 | 0.69 |
| ST3B | 5.83 | 2.99 | 1.25 | 0.61 | 0.01 | 0.64 | 0.78 | 0.81 | 0.83 | 0.03 | 8.135 | 0.58 |
| ST3C | 4.83 | 2.77 | 1.17 | 0.61 | 0.01 | 0.64 | 0.80 | 0.78 | 0.80 | 0.03 | 8.398 | 0.59 |
| ST9 | 5.83 | 3.31 | 1.32 | 0.63 | 0.01 | 0.66 | 0.69 | 0.77 | 0.79 | 0.03 | 9.468 | 0.65 |
| Mean | 6.32 | 2.90 | 1.23 | 0.58 | 0.01 | 0.61 | 0.76 | 0.78 | 0.81 | 0.03 | 8.181 | 0.59 |

Na = Observed number of alleles; Ne = Effective number of alleles; I = Shannon's information statistic; H = Nei's gene diversity; Gst = Genetic differentiation statistics by locus; Uh = Unbiased heterozygosity; Nm = estimate of the number of migrants (gene flow) from Gst where $Nm = 0.5(1 -$ $Gst)/Gst$ ; PIC = Polymorphic information content; Ho = Mean observed heterozygosity over K populations, where $Ho = (\text{Sum (No. of Hets / N)})/k$ ; whereas No. of Hets is number of heterozygotes individuals carrying two different alleles at genetic locus, N is number of individuals in the population and K is number of loci or genetic markers. Hs = Mean expected heterozygosity He over k pops, where $Hs = (\text{Sum } (1 - \text{Sum } p_i^2))/K$ , and $p_i$ = pop allele frequency; Ht = Total expected heterozygosity, where $Ht = 1 - \text{Sum } p_i^2$ , and $p_i$ = average pop allele frequency; F<sub>ST</sub> = Inbreeding coefficient within subpopulations, relative to total genetic differentiation among populations, where $F_{ST} = (Ht - Hs)/Ht$ .

The highest values of Na (10.83), Ne (4.89), H (0.73) and I (1.81) were obtained for locus ST2, while locus ST1 gave the lowest values for these diversity indices (Table 4). The highest (14.7) and lowest (5.13) value for Nm was detected by locus ST6 and ST2, respectively. Except for ST1, which was moderately informative (PIC = 0.42), all the microsatellite loci used were highly informative (PIC >0.5; markers with a PIC value between 0.25 and 0.5 were considered as moderately informative, less than 0.25 as less informative, and more than 0.5 as highly informative).

#### 3.2.1 Genetic variability within and among populations

Average within-population genetic diversity estimates were determined for population sizes ranging from 16 to 76. The populations showed a wide range of within-population genetic differences, with mean values for Ne of 2.89, I of 1.23, H of 0.58, unbiased heterozygosity (Uh) of 0.60, and PPL of 98.61% (Table 5). *Z. tritici* isolates obtained from OSZ showed the highest Na (9.33) value, followed by North Shewa populations. The highest Ne, H, I and Uh values were observed in North Shewa populations, while the lowest Ne and H values were observed in West Shewa populations. The PPL ranged from 91.67% (from West Shewa) to 100% (from OSZ, Arsi, West Arsi and Southwest Shewa) (Table 5).

**Table 5.** Allelic pattern and diversity index for six *Zymoseptoria tritici* populations based on 12 simple sequence repeat (SSR) loci.

| Population | N | Na | Ne | I | H | Uh | PPL |
| --- | --- | --- | --- | --- | --- | --- | --- |
| OSZ | 43 | 9.33 | 3.41 | 1.56 | 0.68 | 0.69 | 100 |
| Arsi | 76 | 7.25 | 2.27 | 1.11 | 0.53 | 0.54 | 100 |
| West Arsi | 26 | 6.67 | 3.04 | 1.33 | 0.62 | 0.64 | 100 |
| SW Shewa | 16 | 3.50 | 2.21 | 0.89 | 0.49 | 0.52 | 100 |
| West Shewa | 18 | 4.33 | 2.16 | 0.93 | 0.47 | 0.50 | 91.67 |
| North Shewa | 21 | 7.27 | 4.25 | 1.58 | 0.69 | 0.72 | 100 |
| Mean | 33.3 | 6.38 | 2.89 | 1.23 | 0.58 | 0.60 | 98.61 |

N = Sample size; Na = Observed number of alleles; Ne = Effective number of alleles; H = Nei's gene diversity; I = Shannon's information index; Uh = Unbiased heterozygosity; PPL = percentage of polymorphic loci. SW Shewa = Southwest Shewa; OSZ = Oromia special zone surrounding Finfinnee.

In the allelic pattern analysis of the 200 isolates, those obtained from Arsi showed the highest number of alleles (9.67), followed by OSZ and North Shewa, which scored 6.83 and 6.33 alleles, respectively. The SWSH population showed the Na (4.08). The highest mean heterozygosity (0.64) was observed in *Z. tritici* from NSH, followed by WSH and Arsi, with mean heterozygosity of 0.62 and 0.59, respectively. The effective number of alleles used for distinguishing populations from each other was highest for the North Shewa population and least (2.21) for the West Arsi population. Likewise, the highest (4.03) number of private alleles was observed in AZ, and the lowest (0.33) in WA and SWSH (Figure 4).

#### 3.2.2 Genetic relationship within and among populations

An AMOVA revealed that 95% of the total variation was attributable to within-population variation, and therefore only 5% to among-population variation (Table 6). The populations exhibited statistically significant but low genetic differentiation (F_ST_ = 0.02; p < 0.001).

**Table 6.** Analysis of molecular variance of 12 simple sequence repeats (SSR) from six populations of *Zymoseptoria tritici*.

| Source of variation | d.f. | Sum of squares | VC | Percentage of variation | FI | p-value |
| --- | --- | --- | --- | --- | --- | --- |
| Among populations | 5 | 50.99 | Va = 0.09 | 2% | FIS = 0.02 | Va and FCT <0.001 |
| Among individuals | 194 | 941.45 | Vb = 0.11 | 3% | FST = 0.02 | Vb and FSC <0.001 |
| Within individuals | 200 | 928.00 | Vc = 4.64 | 95% | FIT = 0.04 | Vc and FST <0.001 |
| Total | 399 | 1920.44 | 4.83 |  |  |  |
| Gene flow (Nm = 14.7) |  |  |  |  |  |  |

Whereas d.f. = degree of freedom, VC = Variance Component and FI = Fixation index, FIT is Overall inbreeding coefficient Calculated as $1 - (HI / HT)$ , FIS is Inbreeding coefficient calculated as $1 - (HI / HS)$ FST = $1 - (HS / HT)$ , Va = variance component among population, Vb = Variance component among individual, Vc = Variance component within individuals.

#### 3.2.3 Genetic relationships between the populations

Table 7 shows the estimates for pairwise genetic distance and gene flow. The highest genetic distances, of 0.30, 0.29 and 0.27, were observed between the populations of North Shewa and Southwest Shewa, North Shewa and Arsi, and North Shewa and West Shewa, respectively. Conversely, the lowest genetic distance (0.06) was observed between the populations of Southwest Shewa and Arsi, and West Shewa and Arsi (Table 7).

**Table 7.**
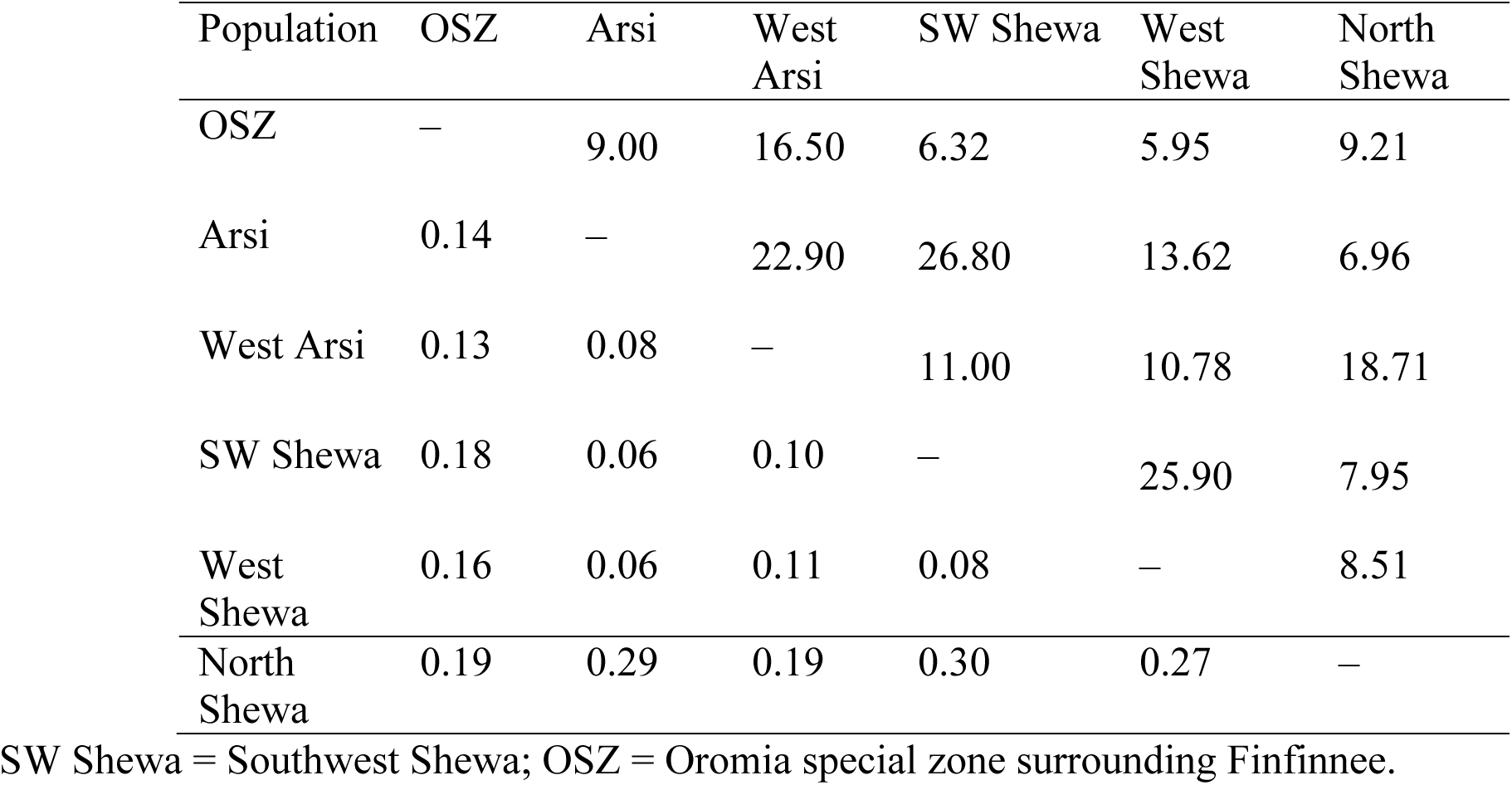
Pairwise Nei’s genetic distance (H) (below diagonal) and gene flow (Nm) (haploid) values (above diagonal) among six *Zymoseptoria tritici* populations from Ethiopia.

The pairwise coefficient of genetic differentiation between the populations ranged from 0.002 (between West Shewa and Southwest Shewa) to 0.077 (between West Shewa and OSZ). The highest and statistically significant genetic differentiation (PhiPT = 0.077, *p* < 0.001), with a gene flow rate of 5.9, was observed between populations of West Shewa and OSZ, implying a relatively low gene flow between them. The second-highest genetic differentiation, with a gene flow rate of 6.33, was observed between the populations of Southwest Shewa and OSZ. The genetic differences between populations of Arsi and West Arsi, Arsi and Southwest Shewa, Southwest Shewa and West Shewa, and West Arsi and North Shewa, were not statistically significant (*p* > 0.05) (Table 8), whereas the corresponding gene flow for these pairs of populations was high (Table 7). The fixation index between the populations was very low, with the highest fixation index (F_ST_ = 0.04) observed between the populations of Southwest Shewa and North Shewa (Figure 5).

**Figure 5.**
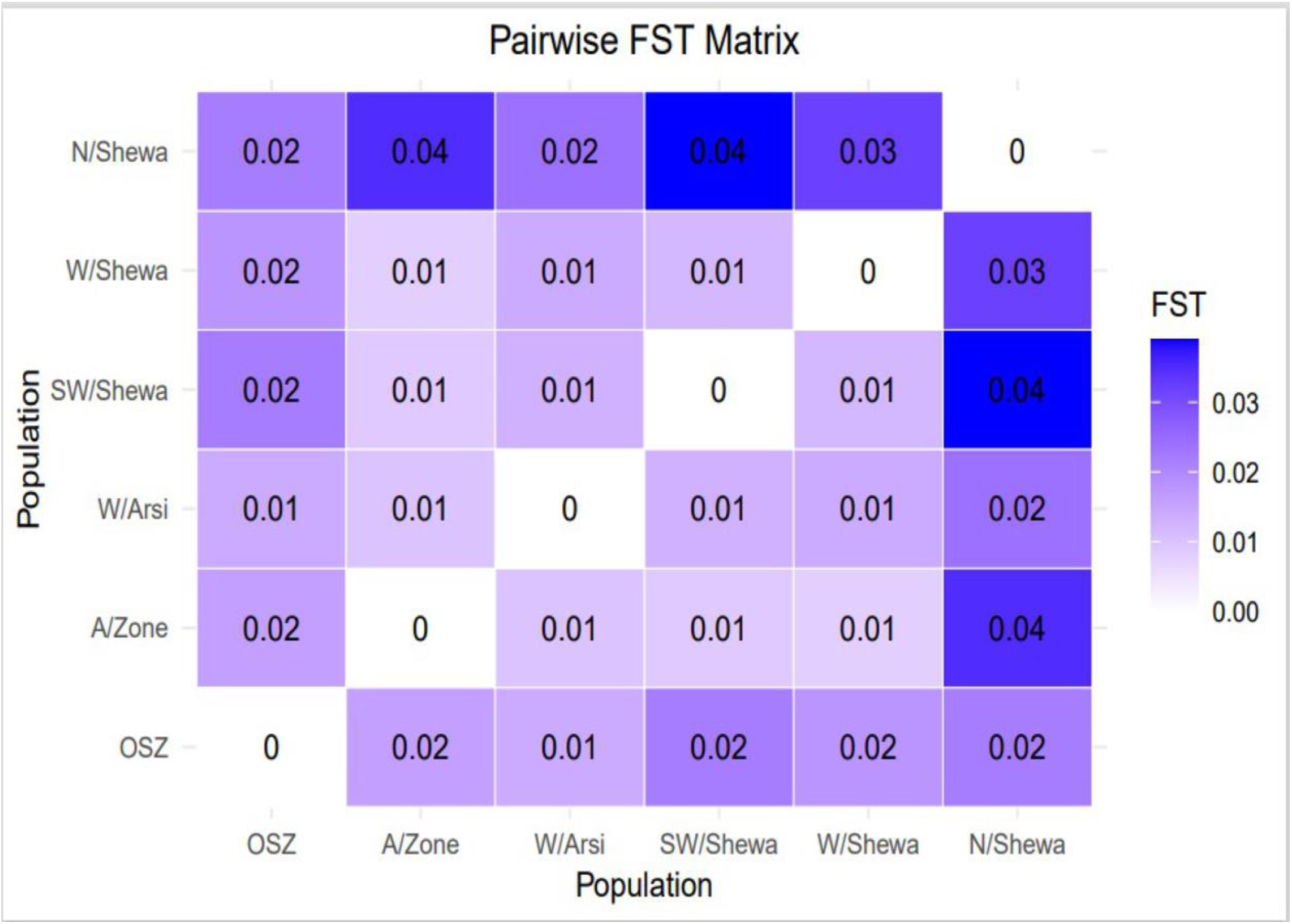
A graphical representation of a pairwise F_ST_ matrix for six populations of *Zymoseptoria tritici*. Whereas OSZ = Oromia special Zone surrounding Finfinne.

**Table 8.** Population genetic differentiation as measured by PhiPT (a genetic differentiation test for the variance between populations/total genetic variants) (below the diagonal) among six *Zymoseptoria tritici* populations from Ethiopia, with *p*-values above the diagonal.

| Population | OSZ | Arsi | West Arsi | SW Shewa | West Shewa | North Shewa |
| --- | --- | --- | --- | --- | --- | --- |
| OSZ | – | 0.001 | 0.004 | 0.001 | 0.001 | 0.001 |
| Arsi | 0.053 | – | 0.010 | 0.057 | 0.001 | 0.001 |
| West Arsi | 0.029 | 0.021 | – | 0.014 | 0.003 | 0.023 |
| SW Shewa | 0.073 | 0.018 | 0.043 | – | 0.402 | 0.003 |
| W/Shewa | 0.077 | 0.035 | 0.044 | 0.002 | – | 0.004 |
| N/Shewa | 0.051 | 0.067 | 0.026 | 0.059 | 0.046 | – |

SW Shewa = South West Shewa; W/Shewa = West Shewa; N/Shewa = North Shewa; OSZ = Oromia special zone surrounding Finfinnee.

#### 3.2.4 Cluster analysis

PCA is a technique frequently used in multivariate statistics to display patterns in genetic structure, and similarly to determine the amounts of variance described per component and cumulatively (Mekonnen *et al*., 2020). In the current study, principal component 1 (PC1) and principal component 2 (PC2) explained 33.51% (PC1 = 20.57% and PC2 =12.94%) of the total genetic variation (Figure 6). PCA clustered the entire population into two subgroups with varying degrees of genetic admixture. The populations of Arsi, West Arsi, North Shewa, West Shewa and Southwest Shewa were clustered together. In contrast, individuals from OSZ populations showed a nearly uniform distribution on the two-dimensional coordinate plane, with little genetic admixture with the other *Z. tritici* population (Figure 6). This finding indicated the presence of a significant gene flow between the geographical areas, resulting in poor clustering of the isolates. None of the clusters was composed entirely of isolates from a particular population (Figure 6).

**Figure 6.**
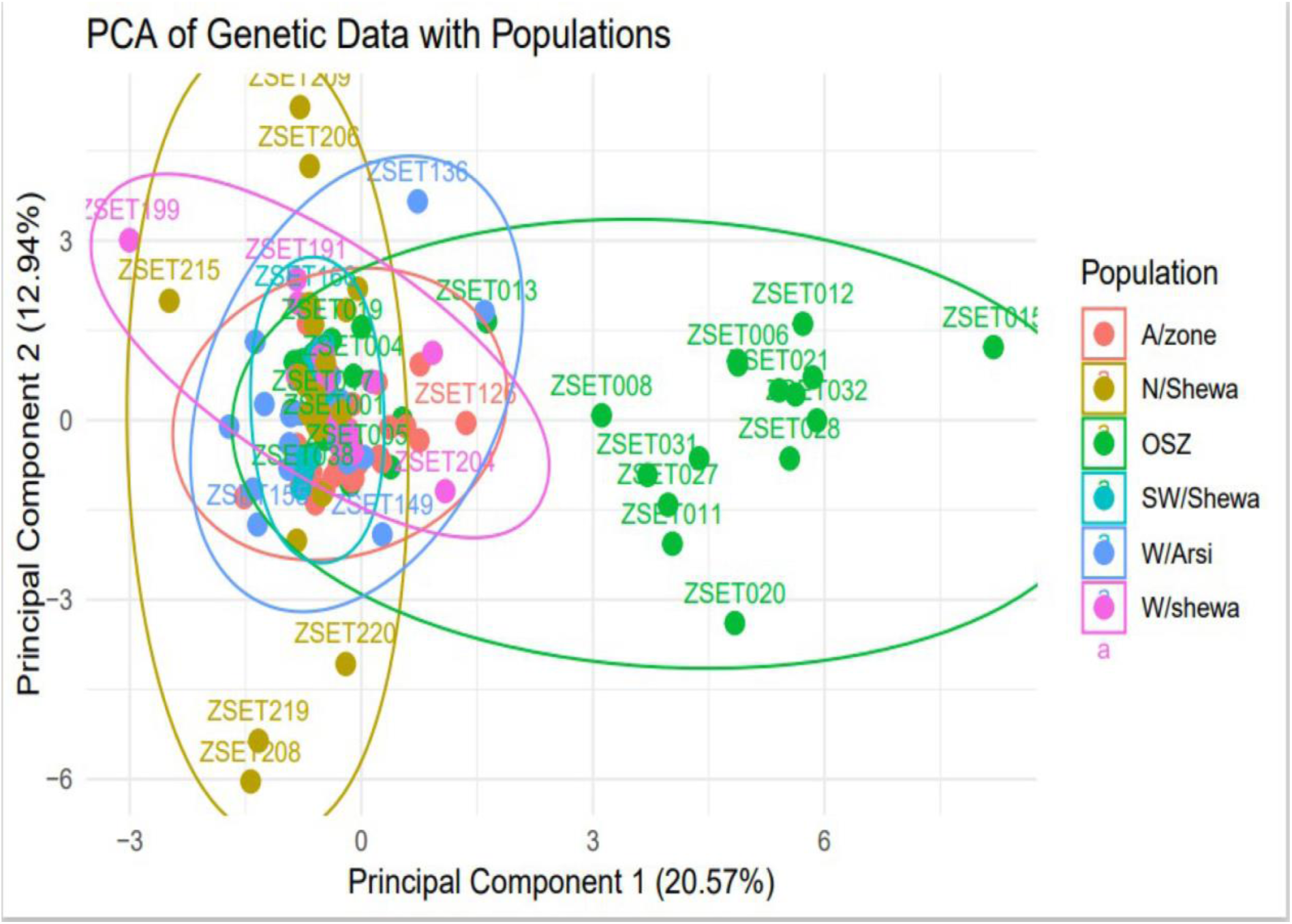
A principal component analysis (PCA) for the 200 individual *Zymoseptoria tritici* isolates as revealed by 12 simple sequence repeat (SSR) markers. Samples coded with the same symbol and color belong to the same population. OSZ = Oromia special Zone.

Accordingly, six populations of *Z. tritici* were sorted into three primary clusters (C1, C2 and C3), each of which was further divided into two sub-clusters based on an unweighted pair group method with arithmetic mean (UPGMA) of the measured of dissimilarity (Figure 7). Of the 200 isolates, 86 individuals (43%) belonged to C1, followed by C2 and C3, each of which had 57 (28.5%) individuals. None of the major clusters was made up of members of a single population, indicating high levels of gene flow. The UPGMA yielded a dendrogram that also divided the six populations into three main clusters (Figure 7).

**Figure 7.**
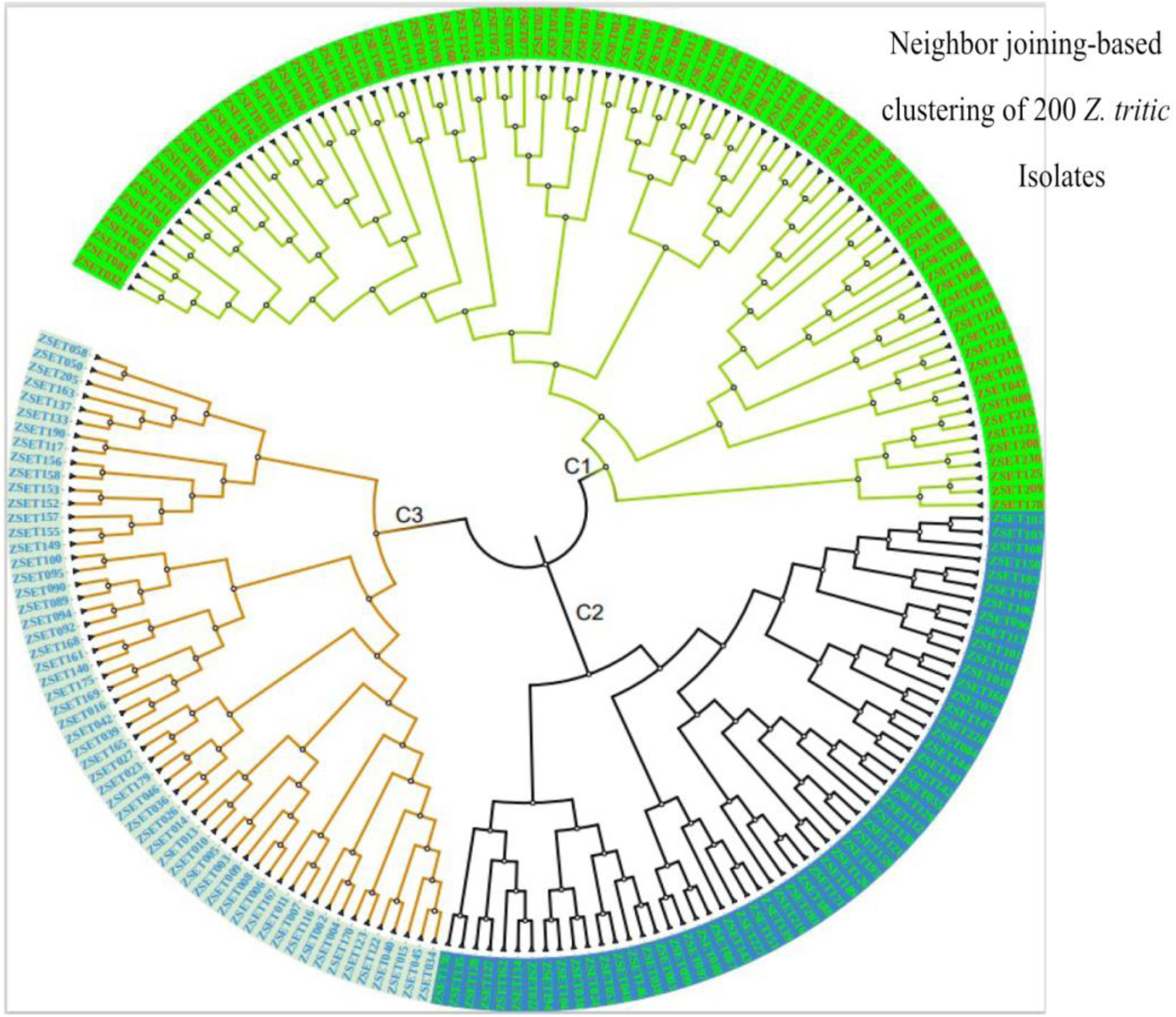
The neighbor joining-based clustering of 200 *Zymoseptoria tritici* isolates representing six populations. Samples coded with the same color belong to the same clade (C1–C3).

#### 3.2.5 Population structure analysis

The population structure analysis identified two genetic groups (K = 2), indicating that the isolates were sourced from two subpopulations (Figure 8). None of the study populations was composed exclusively of isolates from a particular subpopulation. Each isolate shared alleles from both subpopulations, indicating the presence of genetic admixture (Figure 8). Population structure analysis of the 200 *Z. tritici* isolates identified the best delta K value as 2 (Figure 9; the red and green colors represent the proposed isolates the two genetic groups shared).

**Figure 8.**
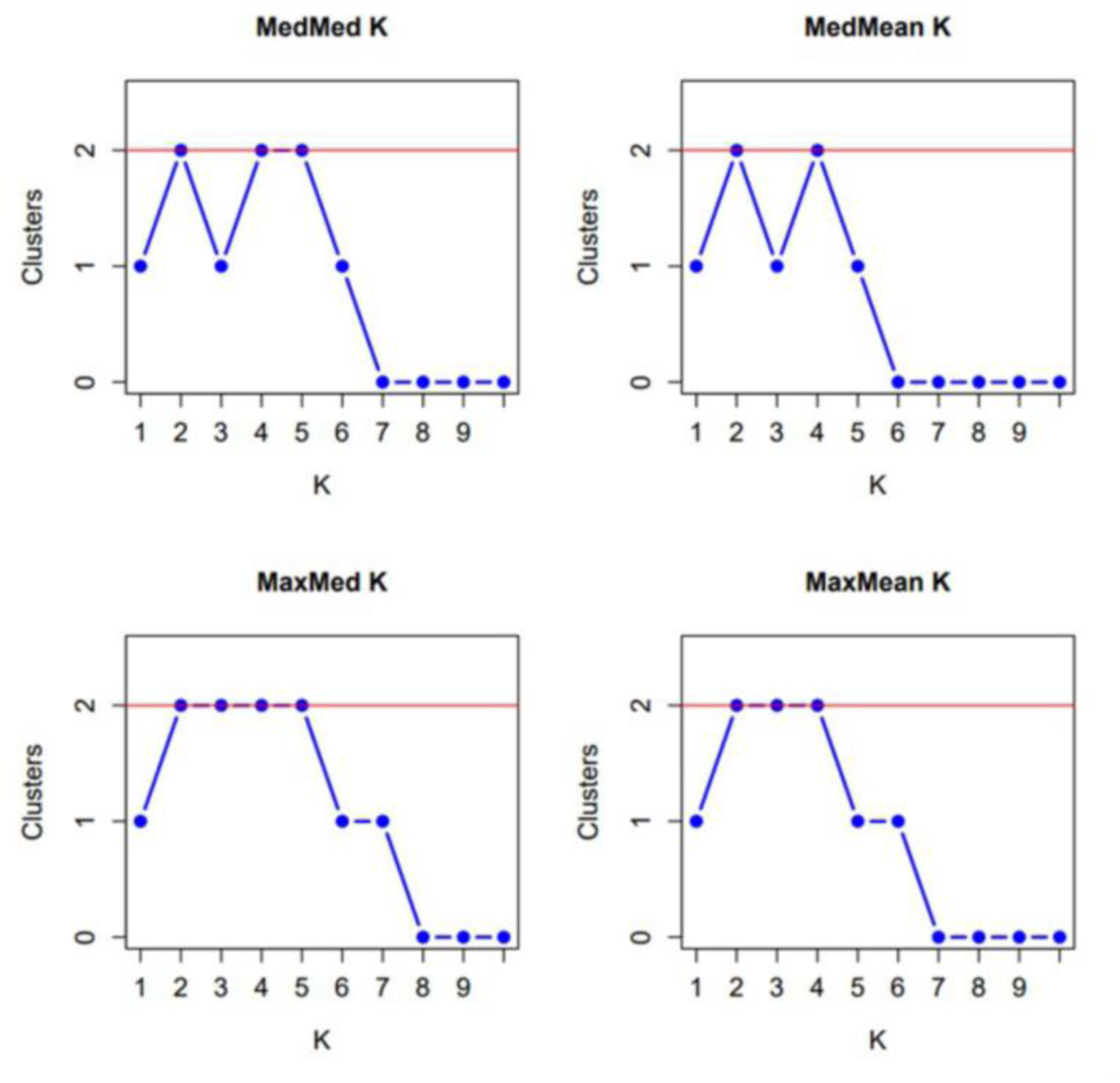
Graphs displaying an optimum of two genetic clusters representing the six *Zymoseptoria tritici* populations based on the approach of (Puechmaille, 2016) where K = Population Cluster (1 - 9).

**Figure 9.**
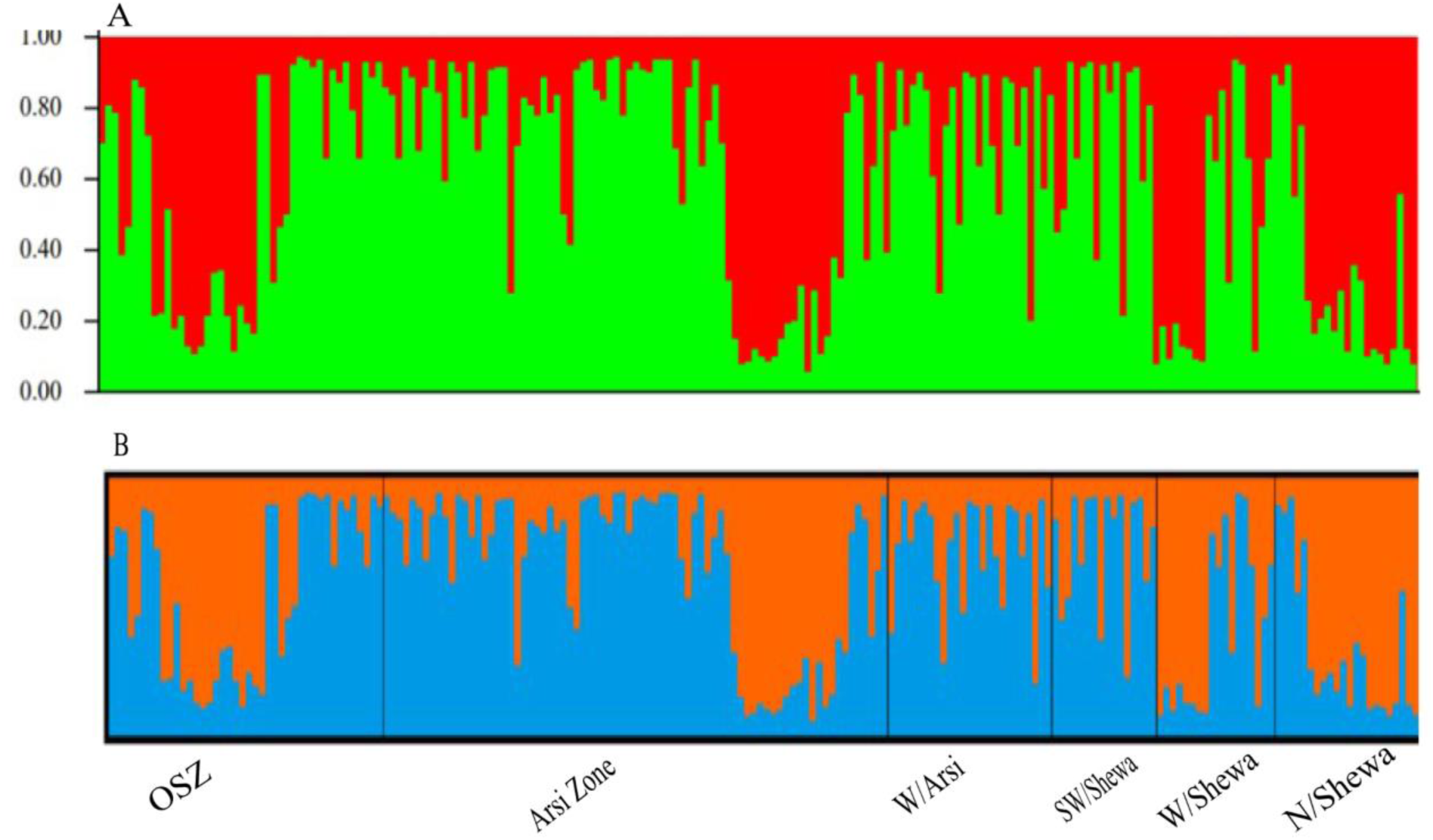
The population genetic structure of 200 individual isolates of *Zymoseptoria tritici* for K = 2 (Figure 8). Each color represents a different cluster, and the different colors of each genotype represent membership in the different genetic populations. **(A)** Graphical representation of individual genotypes arranged according to the level of their membership in different clusters at 65% membership; **(B)** graphical display of the genetic structure of each population [isolates 43, 76, 26, 16, 18 and 21 representing OSZ (Oromia special zone surrounding Finfinnee), Arsi, West Arsi, SW/Shewa (Southwest Shewa) and N/Shewa (North Shewa), respectively].

## 4 Discussion

*Z. tritici* is a haploid organism, and the SSR markers used in this study produced a single allele per isolate at each locus, confirming that they were present as single copies. The most common allele frequency at each locus was less than 0.95 or 0.99, confirming the loci as polymorphic, and thus a useful genetic tool for population genetics studies. There were three to seven alleles per locus, with an average of 5.5 alleles among the 12 loci. The average number of alleles observed in the present study was significantly higher than that reported by (Mekonnen *et al*., 2020), who described an average of 3.5 alleles per locus among 182 isolates of *Z. tritici*, from Ethiopia and the level reported by (Siah *et al*., 2018), who described an average of 4.21 alleles per locus for *Z. tritici* populations of northern France using eight SSR markers.

All the SSR microsatellites used were highly polymorphic and informative (PIC = 0.59), indicating their relevance to unlocking the genetic structure of the target pathogen. They can therefore serve as a useful genetic tool for informing durable and effective management strategies to control STB. The SSR markers used in this study showed a high locus diversity (H = 0.58, Nm =8.18 and PIC 0.59), which is in agreement with the report of (Mekonnen *et al*., 2020), who observed a mean locus genetic diversity of 0.45, gene flow of 2.64 and PIC of 0.49, for 182 *Z. tritici* isolates also from Ethiopia.

The presence of high within-population genetic diversity in the *Z. tritici* populations was confirmed, with mean values for Na, Ne, I, He, H and PPL of 6.32, 2.90, 1.22, 0.58, 0.57 and 98.6%, respectively. The *Z. tritici* populations of Ethiopia appear to have higher within-population diversity than the *Z. tritici* populations of Tunisia, as reported by (Chedli *et al*., 2022), who observed values of Ne = 1.67, I= 0.49, H = 0.28 and %PPL = 62.5 for 162 *Z. tritici* isolates obtained from three regions. Higher genetic diversity has also been reported for *Z. tritici* populations in the United States (Gurung *et al*., 2011), Tunisia (Boukef *et al*., 2012) which was reported as having lower genetic diversity when compared to current study and northern France (El Chartouni *et al*., 2011b) have higher genetic diversity. In the present study, the highest genetic diversity was observed in the *Z. tritici* populations collected from OSZ, followed by North Shewa and then West Arsi and Arsi, confirming these areas as ideal for studies of the pathogen’s genetics and genome, and host–pathogen interactions, and also multi-location wheat germplasm screening for STB resistance.

The AMOVA test revealed the presence of greater genetic diversity (95%) within populations than between populations (5%). The higher within-population variation could be the result of random changes in the sequences of genes in the DNA (mutation) leading to the formation of new genes or alleles, gene movement between the different populations, and sexual reproduction resulting in the formation of new gene combinations. Higher within-population genetic diversity for *Z. tritici* has been reported elsewhere in the world, including in Iran (Mahboubi *et al*., 2022), northern France *(Siah et al*., 2018) and Tunisia (Chedli *et al*., 2022).

The differentiation coefficient Fst is commonly used to assess population structure, with Fst values between 0.00 and 0.05 indicating minimal divergence, and Fst values between 0.05 and 0.15 indicating moderate divergence. The *Z. tritici* populations of OSZ and Southwest Shewa, OSZ and West Shewa, North Shewa and Arsi, and North Shewa and Southwest Shewa, showed moderate divergence (Fst = 0.05–0.15), while all other population pairs showed little population divergence (Fst <0.05).

The *Z. tritici* populations had a level of high gene flow (Nm =14.7), resulting in lower population differentiation (FST = 0.014) (Chedli *et al*., 2022). Gene flow in *Z. tritici* occurs primarily through three mechanisms: the establishment of a sexual reproductive cycle, the existence of intermediate hosts such as some grasses, which can act as a green bridge, and the human transport of infected seeds and straw. The significant level of genetic variability among populations in the current study may be explained by spontaneous mutation and sexual recombination (McDonald and Linde, 2002). Ascospores of *Z. tritici* may move across large distances by wind, which can lead to reduced genetic differentiation between populations (5%). Other factors that might have facilitated high gene flow among isolates from different geographical zones include the movement of plant parts such as straw, the exchange of infected seeds through trading, and long-distance movement of ascospores.

There was no clear, genetically identifiable clustering pattern in the *Z. tritici* populations studied. Neighbor joining-based cluster analyses did not group the populations according to their administrative zones, indicating the presence of close relationships and sharing of alleles among the different *Z. tritici* populations, which probably arose through long-distance movement of the spores by air and/or germplasm exchanges in the form of seeds from common market places. *Z. tritici* populations from different administrative zones showed significant genetic admixture, resulting in high gene flow, and hence PCA and STRUCTURE analyses were also unable to cluster the populations effectively based on the geographic distribution of the sampling areas. A Bayesian model-based clustering algorithm was used to determine the number of subpopulations (K) given an admixture model with associated allele frequencies. The STRUCTURE analysis supported the result of the PCA, and indicated that the *Z. tritici* populations were sourced from two (K = 2) subpopulations, with a greater degree of genetic admixture among the *Z. tritici* isolates of the different administrative zones as a result of high gene flow. Similar population genetic admixtures have been documented for *Z. tritici* populations in Tunisia (Chedli *et al*., 2022).

## 5 Conclusion

STB caused by *Z. tritici* (*M. graminicola*) represents a serious threat to global wheat production, and is a major bottleneck to wheat production in Ethiopia, causing significant yield loss. In order to develop and implement long-lasting and successful management strategies, accurate identification and data on the genetic makeup of pathogen populations gathered from hotspot sites are crucial. In this study, a total of 167 *Z. tritici* isolates was successfully characterized by sequencing ITS rDNA regions. Sequence-based phylogenetic analyses revealed a high level of genetic admixture among the *Z. tritici* isolates collected from various administrative zones. The genetic structure analysis of 200 *Z. tritici* isolates recovered from six administrative zones using 12 SSR revealed high levels of genetic diversity, with 95% of the total genetic variation residing within populations, confirming that sexual recombination is taking place at high levels in the *Z. tritici* populations of central and south-eastern Ethiopia. Among the six populations, those obtained from OSZ, Arsi, West Arsi and North Shewa showed the highest genetic diversity. These locations therefore provide hotspots for *Z. tritici* genetic analyses and can serve as good environments for screening wheat germplasm for resistance to STB. The *Z. tritici* populations exhibited low to moderate pairwise genetic differentiation (Fst = 0.05–0.15), probably because of the high gene flow rate (Nm = 14.7). Because of the significant gene flow in the pathogen populations caused by long-distance spore movement and the exchange of infected seeds through trading, clustering, PCA and STRUCTURE analyses were unable to separate the populations based on their sampling areas. To confirm the present results, and to gain further insight into the pathogen’s genetic structure, we recommend continued analyses of pathogen populations, including from other wheat major wheat-growing areas, as well as populations obtained from different years at different growth stages of the pathogen’s preferred (both bread and durum wheat) and alternative hosts.

## Supporting information

All Supplementary tables and figures

## Acknowledgments

The authors extend their heartfelt gratitude to the wheat farming community in the study areas for their kind permission to assess their fields and sample STB-symptomatic wheat leaves. We are also deeply thankful to the National Agricultural Biotechnology Research Center (NABRC) in Holeta, Ethiopia, for providing the laboratory space and facilities essential for this research. Similarly, our appreciation goes to the Institute of Biotechnology at Addis Ababa University for their technical support throughout the study. Furthermore, RRV acknowledge support from FORMAS (2019–01316), Carl Tryggers Stiftelse för Vetenskaplig Forskning (CTS 20: 464), NOVO Nordisk Foundation (0074727), and SLU Centre for Biological Control.

## Notes

### Competing Interest Statement

The authors have declared no competing interest.

