## Supplementary material for "Genetic Diversity of *Zymoseptoria tritici* Populations in Central and South-eastern Ethiopia": All Supplementary tables and figures

Supplementary Table 1. Global Positioning System (GPS) data showing the six administrative zones covered for *Z. tritici* isolates collection.

| NO | Sample ID | Population | Latitude (X -axis) | Altitude (Y-axis) |
|---|---|---|---|---|
| 1 | ZSET001 | OSZ | 9.0569269 | 38.5069977 |
| 2 | ZSET002 | OSZ | 9.0597527 | 38.5068122 |
| 3 | ZSET003 | OSZ | 9.0597444 | 38.5068011 |
| 4 | ZSET004 | OSZ | 9.0596175 | 38.5068292 |
| 5 | ZSET005 | OSZ | 9.0596485 | 38.5068793 |
| 6 | ZSET006 | OSZ | 9.0595688 | 38.5071029 |
| 7 | ZSET007 | OSZ | 9.0596034 | 38.5071266 |
| 8 | ZSET008 | OSZ | 9.0596284 | 38.5071872 |
| 9 | ZSET009 | OSZ | 9.0596105 | 38.5071828 |
| 10 | ZSET010 | OSZ | 9.0594488 | 38.5072152 |
| 11 | ZSET011 | OSZ | 9.0594472 | 38.5072374 |
| 12 | ZSET012 | OSZ | 9.0594485 | 38.5072549 |
| 13 | ZSET013 | OSZ | 9.059435 | 38.5072471 |
| 14 | ZSET014 | OSZ | 9.0594342 | 38.5072336 |
| 15 | ZSET015 | OSZ | 9.0603669 | 38.5073142 |
| 16 | ZSET016 | OSZ | 9.0603775 | 38.5073718 |
| 17 | ZSET017 | OSZ | 9.0605835 | 38.5075603 |
| 18 | ZSET018 | OSZ | 9.0603903 | 38.507583 |
| 19 | ZSET019 | OSZ | 9.0605595 | 38.5075624 |
| 20 | ZSET020 | OSZ | 9.0605604 | 38.5076252 |
| 21 | ZSET021 | OSZ | 9.0605696 | 38.5077458 |
| 22 | ZSET022 | OSZ | 9.0605607 | 38.5077651 |
| 23 | ZSET023 | OSZ | 9.0605514 | 38.5077176 |
| 24 | ZSET024 | OSZ | 9.060557 | 38.5077263 |
| 25 | ZSET025 | OSZ | 9.0604949 | 38.5078732 |
| 26 | ZSET026 | OSZ | 9.0569553 | 38.507404 |
| 27 | ZSET027 | OSZ | 9.0569129 | 38.5073957 |
| 28 | ZSET028 | OSZ | 9.0569022 | 38.5075294 |
| 29 | ZSET029 | OSZ | 9.056879 | 38.5074794 |
| 30 | ZSET030 | OSZ | 9.056807 | 38.5074122 |
| 31 | ZSET031 | OSZ | 9.0567543 | 38.5074043 |
| 32 | ZSET032 | OSZ | 9.05654 | 38.5072454 |
| 33 | ZSET033 | OSZ | 9.0565292 | 38.5072762 |

| | | | | |
|---|---|---|---|---|
| 34 | ZSET034 | OSZ | 9.0563627 | 38.5071617 |
| 35 | ZSET035 | OSZ | 9.056241 | 38.5071142 |
| 36 | ZSET036 | OSZ | 9.0562057 | 38.5070435 |
| 37 | ZSET037 | OSZ | 9.0564594 | 38.5070058 |
| 38 | ZSET038 | OSZ | 9.0564453 | 38.507015 |
| 39 | ZSET039 | OSZ | 8.9273379 | 38.8190416 |
| 40 | ZSET040 | OSZ | 8.9273469 | 38.8190094 |
| 41 | ZSET041 | OSZ | 8.9273442 | 38.8189631 |
| 42 | ZSET042 | OSZ | 8.8809197 | 38.8163993 |
| 43 | ZSET043 | OSZ | 9.0569269 | 38.5069977 |
| 44 | ZSET044 | A/zone | 8.1747366 | 39.2432836 |
| 45 | ZSET045 | A/zone | 8.1487019 | 39.2368839 |
| 46 | ZSET046 | A/zone | 8.1487761 | 39.2367133 |
| 47 | ZSET047 | A/zone | 8.1085819 | 39.2282024 |
| 48 | ZSET047 | A/zone | 8.064613 | 39.2173333 |
| 49 | ZSET048 | A/zone | 8.0415884 | 39.1970149 |
| 50 | ZSET049 | A/zone | 8.0201173 | 39.1571817 |
| 51 | ZSET050 | A/zone | 8.0201061 | 39.1571715 |
| 52 | ZSET052 | A/zone | 8.0203317 | 39.1559175 |
| 53 | ZSET053 | A/zone | 8.0207988 | 39.1558163 |
| 54 | ZSET054 | A/zone | 8.0206275 | 39.155814 |
| 55 | ZSET055 | A/zone | 8.0207357 | 39.1557938 |
| 56 | ZSET057 | A/zone | 8.0205833 | 39.1558425 |
| 57 | ZSET058 | A/zone | 8.0211456 | 39.1563795 |
| 58 | ZSET060 | A/zone | 8.0205748 | 39.1561334 |
| 59 | ZSET061 | A/zone | 8.0216847 | 39.1547277 |
| 60 | ZSET062 | A/zone | 8.0212884 | 39.154875 |
| 61 | ZSET064 | A/zone | 8.0217344 | 39.1538325 |
| 62 | ZSET065 | A/zone | 8.0207542 | 39.1550355 |
| 63 | ZSET067 | A/zone | 8.0220362 | 39.1546981 |
| 64 | ZSET070 | A/zone | 8.0209196 | 39.154974 |
| 65 | ZSET072 | A/zone | 8.0211844 | 39.1546004 |
| 66 | ZSET073 | A/zone | 8.0208715 | 39.1549649 |
| 67 | ZSET074 | A/zone | 8.0209997 | 39.1549862 |
| 68 | ZSET075 | A/zone | 8.0215224 | 39.1546602 |
| 69 | ZSET076 | A/zone | 8.0211649 | 39.1546203 |
| 70 | ZSET077 | A/zone | 7.9571685 | 39.1265004 |
| 71 | ZSET079 | A/zone | 7.8810894 | 39.1246496 |
| 72 | ZSET079 | A/zone | 7.88106 | 39.124785 |

| | | | | |
|---|---|---|---|---|
| 73 | ZSET080 | A/zone | 7.8810691 | 39.1249253 |
| 74 | ZSET082 | A/zone | 7.8810242 | 39.1251437 |
| 75 | ZSET083 | A/zone | 7.8808577 | 39.1251743 |
| 76 | ZSET084 | A/zone | 7.8427269 | 39.1380444 |
| 77 | ZSET085 | A/zone | 7.8422822 | 39.138437 |
| 78 | ZSET086 | A/zone | 7.8028474 | 39.1424028 |
| 79 | ZSET087 | A/zone | 7.8028749 | 39.142656 |
| 80 | ZSET088 | A/zone | 7.8025738 | 39.1427628 |
| 81 | ZSET089 | A/zone | 7.8025157 | 39.1427599 |
| 82 | ZSET090 | A/zone | 7.7912048 | 39.1488327 |
| 83 | ZSET091 | A/zone | 7.7912829 | 39.148794 |
| 84 | ZSET092 | A/zone | 7.7912544 | 39.1488595 |
| 85 | ZSET094 | A/zone | 7.6295044 | 39.224966 |
| 86 | ZSET095 | A/zone | 7.6294857 | 39.2249666 |
| 87 | ZSET096 | A/zone | 7.5998192 | 39.230454 |
| 88 | ZSET097 | A/zone | 7.5998809 | 39.2304643 |
| 89 | ZSET098 | A/zone | 7.5997786 | 39.2303337 |
| 90 | ZSET099 | A/zone | 7.5997209 | 39.2304392 |
| 91 | ZSET100 | A/zone | 7.5797133 | 39.2431636 |
| 92 | ZSET101 | A/zone | 7.5797652 | 39.2431495 |
| 93 | ZSET102 | A/zone | 7.5799478 | 39.2431251 |
| 94 | ZSET103 | A/zone | 7.5799521 | 39.2432951 |
| 95 | ZSET104 | A/zone | 7.5437801 | 39.2552885 |
| 96 | ZSET105 | A/zone | 7.5438331 | 39.2552222 |
| 97 | ZSET106 | A/zone | 7.5442644 | 39.255244 |
| 98 | ZSET107 | A/zone | 7.5442459 | 39.255319 |
| 99 | ZSET108 | A/zone | 7.5446436 | 39.25584 |
| 100 | ZSET109 | A/zone | 7.5446458 | 39.2558506 |
| 101 | ZSET110 | A/zone | 7.544628 | 39.2560705 |
| 102 | ZSET112 | A/zone | 7.5448962 | 39.2560249 |
| 103 | ZSET113 | A/zone | 7.5447388 | 39.2557504 |
| 104 | ZSET114 | A/zone | 7.5447398 | 39.255736 |
| 105 | ZSET115 | A/zone | 7.5446688 | 39.2557108 |
| 106 | ZSET116 | A/zone | 7.5446858 | 39.2557205 |
| 107 | ZSET117 | A/zone | 7.5446853 | 39.2556933 |
| 108 | ZSET118 | A/zone | 7.5446774 | 39.2556623 |
| 109 | ZSET119 | A/zone | 7.5446602 | 39.2556463 |
| 110 | ZSET120 | A/zone | 7.5444003 | 39.255155 |
| 111 | ZSET121 | A/zone | 7.5444045 | 39.2551809 |

| | | | | |
|---|---|---|---|---|
| 112 | ZSET122 | A/zone | 7.5443574 | 39.2551551 |
| 113 | ZSET123 | A/zone | 7.5443482 | 39.255141 |
| 114 | ZSET124 | A/zone | 7.5443438 | 39.255157 |
| 115 | ZSET125 | A/zone | 7.5443406 | 39.2551559 |
| 116 | ZSET126 | A/zone | 7.5443333 | 39.2551352 |
| 117 | ZSET127 | A/zone | 7.5441266 | 39.2550439 |
| 118 | ZSET128 | A/zone | 7.5441097 | 39.2550609 |
| 119 | ZSET129 | A/zone | 7.4777469 | 39.2620539 |
| 120 | ZSET130 | W/Arsi | 7.3715682 | 39.2542695 |
| 121 | ZSET131 | W/Arsi | 7.3714953 | 39.2543007 |
| 122 | ZSET132 | W/Arsi | 7.0837071 | 38.7862882 |
| 123 | ZSET133 | W/Arsi | 7.0837347 | 38.7863517 |
| 124 | ZSET134 | W/Arsi | 7.0834679 | 38.786532 |
| 125 | ZSET136 | W/Arsi | 7.083497 | 38.7865181 |
| 126 | ZSET137 | W/Arsi | 7.0835245 | 38.7865191 |
| 127 | ZSET138 | W/Arsi | 7.0835192 | 38.7865106 |
| 128 | ZSET139 | W/Arsi | 7.0835322 | 38.7865187 |
| 129 | ZSET140 | W/Arsi | 7.0835361 | 38.7865125 |
| 130 | ZSET141 | W/Arsi | 7.0835375 | 38.7865091 |
| 131 | ZSET142 | W/Arsi | 7.0835409 | 38.7865042 |
| 132 | ZSET143 | W/Arsi | 7.0839987 | 38.786236 |
| 133 | ZSET144 | W/Arsi | 7.0840209 | 38.7862003 |
| 134 | ZSET145 | W/Arsi | 7.0842813 | 38.7861392 |
| 135 | ZSET146 | W/Arsi | 7.0843064 | 38.7861087 |
| 136 | ZSET147 | W/Arsi | 7.0843233 | 38.7861251 |
| 137 | ZSET148 | W/Arsi | 7.0840605 | 38.7863886 |
| 138 | ZSET149 | W/Arsi | 7.0840369 | 38.7865058 |
| 139 | ZSET150 | W/Arsi | 7.0840395 | 38.7865294 |
| 140 | ZSET151 | W/Arsi | 7.0840259 | 38.7865139 |
| 141 | ZSET152 | W/Arsi | 7.084013 | 38.7865099 |
| 142 | ZSET153 | W/Arsi | 7.0838573 | 38.7865496 |
| 143 | ZSET155 | W/Arsi | 7.0843883 | 38.7872663 |
| 144 | ZSET156 | W/Arsi | 7.0202747 | 38.9988112 |
| 145 | ZSET157 | W/Arsi | 7.014381 | 39.0288142 |
| 146 | ZSET158 | SW/Shewa | 8.1224615 | 39.2874394 |
| 147 | ZSET159 | SW/Shewa | 8.1225958 | 39.2873934 |
| 148 | ZSET160 | SW/Shewa | 8.6891972 | 38.2384404 |
| 149 | ZSET161 | SW/Shewa | 8.6323209 | 38.0417685 |
| 150 | ZSET162 | SW/Shewa | 8.6323779 | 38.0416194 |

| | | | | |
|---|---|---|---|---|
| 151 | ZSET163 | SW/Shewa | 8.6323652 | 38.0417241 |
| 152 | ZSET165 | SW/Shewa | 8.6323268 | 38.041489 |
| 153 | ZSET166 | SW/Shewa | 8.6323252 | 38.0414472 |
| 154 | ZSET167 | SW/Shewa | 8.6323454 | 38.0414858 |
| 155 | ZSET168 | SW/Shewa | 8.6324223 | 38.0414516 |
| 156 | ZSET169 | SW/Shewa | 8.6324054 | 38.0414585 |
| 157 | ZSET170 | SW/Shewa | 8.6318198 | 38.0413837 |
| 158 | ZSET171 | SW/Shewa | 8.6172886 | 38.0316986 |
| 159 | ZSET175 | SW/Shewa | 8.6173279 | 38.0317713 |
| 160 | ZSET178 | SW/Shewa | 8.6173265 | 38.0317812 |
| 161 | ZSET179 | SW/Shewa | 8.6173707 | 38.031784 |
| 162 | ZSET188 | W/shewa | 8.6598594 | 37.8911229 |
| 163 | ZSET189 | W/shewa | 8.6737157 | 37.8838739 |
| 164 | ZSET190 | W/shewa | 8.673814 | 37.8838156 |
| 165 | ZSET191 | W/shewa | 8.6736658 | 37.8837858 |
| 166 | ZSET192 | W/shewa | 8.6799812 | 37.8882704 |
| 167 | ZSET193 | W/shewa | 8.6798205 | 37.8882773 |
| 168 | ZSET194 | W/shewa | 8.6798384 | 37.8883107 |
| 169 | ZSET195 | W/shewa | 8.6801966 | 37.8881866 |
| 170 | ZSET196 | W/shewa | 8.7470752 | 37.8791774 |
| 171 | ZSET197 | W/shewa | 8.8105019 | 37.874691 |
| 172 | ZSET198 | W/shewa | 8.8721521 | 37.8924064 |
| 173 | ZSET199 | W/shewa | 8.8722338 | 37.8922632 |
| 174 | ZSET200 | W/shewa | 8.8722454 | 37.8921219 |
| 175 | ZSET201 | W/shewa | 8.8722 | 37.8921166 |
| 176 | ZSET202 | W/shewa | 8.8855991 | 37.8889731 |
| 177 | ZSET203 | W/shewa | 8.8978021 | 37.8840987 |
| 178 | ZSET204 | W/shewa | 8.8986769 | 37.8834042 |
| 179 | ZSET205 | W/shewa | 8.8983834 | 37.8832602 |
| 180 | ZSET206 | N/Shewa | 9.2336907 | 38.7591871 |
| 181 | ZSET207 | N/Shewa | 9.2336843 | 38.7593731 |
| 182 | ZSET208 | N/Shewa | 9.3732933 | 38.7861876 |
| 183 | ZSET209 | N/Shewa | 9.5876111 | 38.8610995 |
| 184 | ZSET210 | N/Shewa | 9.5876923 | 38.8611255 |
| 185 | ZSET211 | N/Shewa | 9.5876248 | 38.8612242 |
| 186 | ZSET212 | N/Shewa | 9.7200233 | 38.8189879 |
| 187 | ZSET213 | N/Shewa | 9.8121388 | 38.5577201 |
| 188 | ZSET214 | N/Shewa | 9.8119593 | 38.5576232 |
| 189 | ZSET215 | N/Shewa | 9.8118599 | 38.5578467 |

| | | | | |
|---|---|---|---|---|
| 190 | ZSET216 | N/Shewa | 9.8118214 | 38.5579527 |
| 191 | ZSET218 | N/Shewa | 9.8117915 | 38.5579083 |
| 192 | ZSET219 | N/Shewa | 9.807808 | 38.54955 |
| 193 | ZSET220 | N/Shewa | 9.8077031 | 38.5495874 |
| 194 | ZSET221 | N/Shewa | 9.8076176 | 38.5494552 |
| 195 | ZSET224 | N/Shewa | 9.8076429 | 38.5493765 |
| 196 | ZSET225 | N/Shewa | 9.8012966 | 38.5456616 |
| 197 | ZSET226 | N/Shewa | 9.8014966 | 38.5458269 |
| 198 | ZSET227 | N/Shewa | 9.7976163 | 38.5428452 |
| 199 | ZSET228 | N/Shewa | 9.7976281 | 38.5427114 |
| 200 | ZSET230 | N/Shewa | 9.7978007 | 38.5424305 |

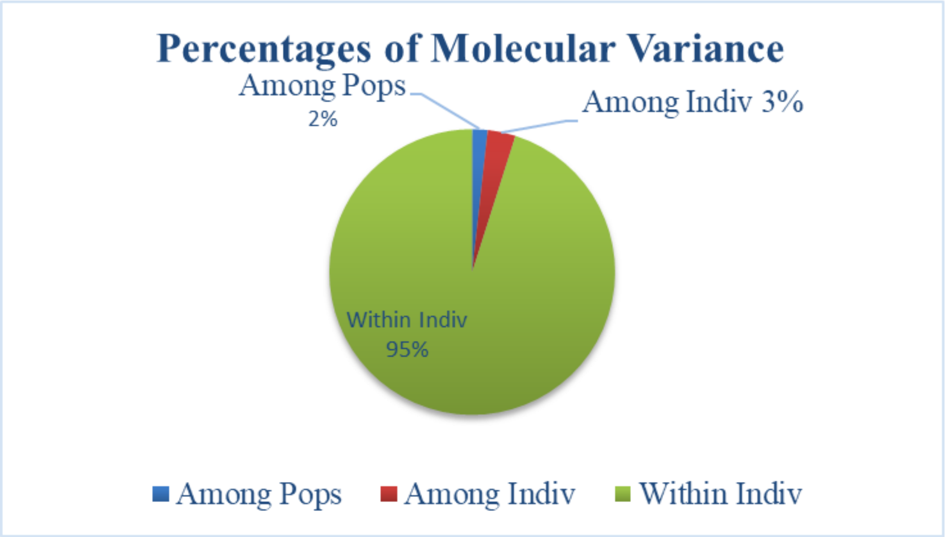


Supplementary Figure 1. Percentage of molecular variance among population and within population.

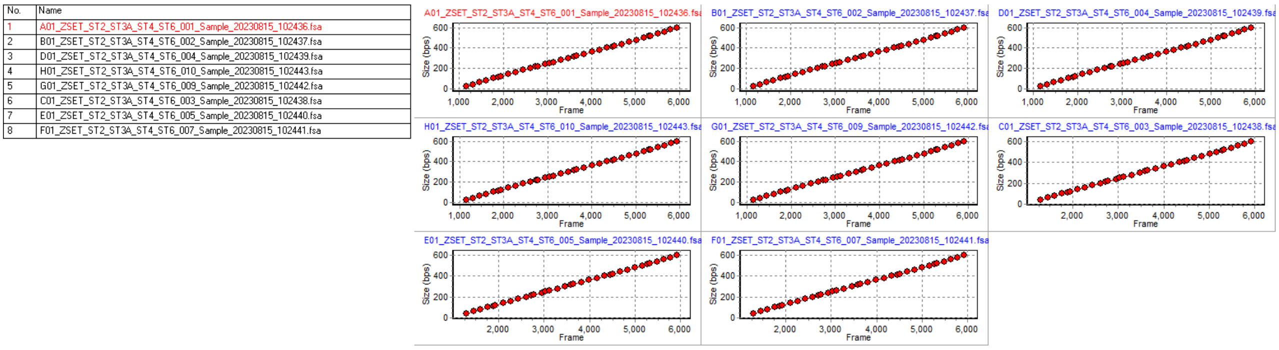

Supplementary Figure 2. Calibration chart of the sample during capillary electrophoresis.
